# Spatial transcriptomics maps molecular and cellular requirements for CD4^+^ T cell-dependent immunity to malaria

**DOI:** 10.1101/2023.02.23.529309

**Authors:** Cameron G. Williams, Marcela L. Moreira, Takahiro Asatsuma, Oliver P. Skinner, Hyun Jae Lee, Shihan Li, Irving Barrera, Evan Murray, Megan S. F. Soon, Jessica A. Engel, David S. Khoury, Saba Asad, Thiago Mass Steiner, Rainon Joseph, Yannick Alexandre, Scott N. Mueller, Fei Chen, Ashraful Haque

## Abstract

CD4^+^ T cells orchestrate adaptive immunity to circulating malaria parasites; yet cellular interactions and molecular mechanisms controlling Th1 and Tfh differentiation in the spleen remain to be fully defined *in vivo*. Here, using a murine model of CD4-dependent immunity, we tested if *Slide-seqV2*, a spatial transcriptomic method with near single-cell resolution, could determine the locations of multiple CD4^+^ T cell subsets and potentially interacting cellular partners in the spleen during infection. Firstly, *Slide-seqV2* readily mapped splenic cellular structure and microanatomical change during infection. Next, computational integration with scRNA-seq reference datasets of splenocytes, stromal cells, and specifically of polyclonal CD4^+^ T cells and B cells, mapped the relative locations of multiple cell-types within this dense tissue. scRNA-seq of B cells over time mapped emergence of germinal centre B cells, red pulp-located plasmablasts and atypical B cells, and uncovered a prolonged CD4^+^ T-cell-independent, follicular bystander B cell response marked by Sca-1 and Ly6C upregulation. scRNA-seq of activated, polyclonal CD4^+^ T cells revealed their similarity to our previous TCR transgenic models. Importantly, spatial analysis revealed polyclonal Th1 cells co-localised with CXCL9/10-producing monocytes in the red pulp, while polyclonal Tfh-like cells were located close to CXCL13-expressing B cell follicles, consistent with our previous CXCR3/CXCR5 competition model of Th1/Tfh bifurcation. CRISPR/Cas9 disruption of either or both CXCR3 and CXCR5 in naïve *Plasmodium*-specific CD4^+^ T cells had unexpectedly minor effects on Th1 differentiation *in vivo*. Instead, CXCR5 was essential for maximising clonal expansion, suggesting a role for splenic CXCL13^+^ cells in supporting CD4^+^ T cell proliferation in malaria. Thus, spatial transcriptomics at near single-cell resolution was feasible in densely packed secondary lymphoid tissue, providing multiple insights into mechanisms controlling splenic polyclonal CD4^+^ T cell and B cell differentiation during infection.

**Highlights:**

- *Slide-seqV2* maps splenic microanatomy, including stromal and immune cell location.
- Bystander activation of all follicular B cells occurs in malaria, marked by Sca-1/Ly6C upregulation.
- Single naïve polyclonal CD4^+^ T cells differentiate mostly into Th1 and Tfh cells in malaria.
- Cell-cell colocalization analysis positions Th1 cells with monocytes in red pulp, and Tfh cells with *Cxcl13^+^* B cell follicles.
- CXCR5, but not CXCR3, supports parasite-specific CD4^+^ T cell clonal expansion.

## Introduction

Naturally-acquired adaptive immunity to malaria in humans often requires multiple exposures over several months to years [1]. Such immunity can wane and is not yet reliably induced by licensed vaccines. A better understanding of how protective cellular and humoral immunity is generated in the spleen during blood-stage *Plasmodium* infection, for example using experimental animal models, may offer strategies for improving control of malaria.

CD4^+^ TCRαβ^+^ T cells and B cells orchestrate cellular and humoral immunity to many microbial pathogens, including *Plasmodium* parasites [2, 3]. To assist in clearing *Plasmodium* parasites as they circulate and replicate in red blood cells, naïve parasite-specific CD4^+^ T cells first differentiate in the spleen into either or both T helper 1 cells (Th1) [4] and T follicular helper cells (Tfh) [5–7]. Th1 cells secrete IFNγ, which is thought to increase phagocytic activity of macrophages [8], and influence immunoglobulin isotype class-switching in B cells [9, 10]. Tfh cells select for affinity-matured B cells within the germinal centre, thus promoting antibody-mediated immunity [11, 12]. CD4^+^ T cells also provide critical ICOS- and CD40L-dependent early assistance to B cells [13–15]. Naïve B cells differentiate into various states including plasmablasts, germinal centre (GC) B cells, memory B cells, atypical B cells, and long-lived plasma cells (LLPC) in malaria [16, 17], with evidence of both parasite-specific IgG and IgM providing protection [18, 19]. Some CD4^+^ T cell and B cell responses may be suboptimal, with plasmablasts potentially acting as an early nutrient sink [20], Th1-polarized Tfh cells failing to assist B cells [21, 22], and regulatory T cells (Tregs) and atypical memory B cells (atMBCs) serving to limit adaptive immunity [23, 24]. Thus, the landscape of CD4^+^ T cell and B cell differentiation within the spleen appears to consist of numerous protective and potentially detrimental cell-states emerging simultaneously. These have so far eluded systematic, genome-scale assessment.

Single-cell mRNA-sequencing (scRNA-seq) has facilitated genome-scale assessments of immune cells in numerous contexts, usually after their recovery as viable cells from blood or tissues of mice and humans [25, 26]. More recently, spatial transcriptomics platforms serve to contextualise transcriptomic data in its tissue location [27–30]. To perform untargeted spatial assessments of mRNA transcripts at near single-cell resolution, we previously developed *SlideseqV2*, which locally captured hundreds to thousands of unique polyadenylated mRNA sequences per positionally barcoded bead on circular arrays comprised of tens of thousands of beads [31, 32]. Although this was validated on murine brain, kidney, and liver tissue, whether *Slide-seqV2* could resolve large numbers of different immune cell-types and activation states, as well as stromal cells, within a densely packed and dynamic secondary lymphoid organ had remained untested.

Previously, we used scRNA-seq to map differentiation of TCR-transgenic CD4^+^ T cells, termed PbTII cells (with specificity for a single epitope in *Plasmodium* Heat Shock Protein 90), during experimental malaria in mice [7, 15, 33]. Single naïve PbTII cells differentiated into Th1 and Tfh-like states during the first week of infection, with cellular depletion studies implicating CXCL9/10^+^ monocytes and B cells in controlling Th1/Tfh fate bifurcation. However, evidence of monocytes and B cells physically interacting with Th1, Tfh, and other CD4^+^ T cell-states has been lacking, particularly for TCR diverse polyclonal cells. Moreover, although early co-expression of CXCR3 and CXCR5 by PbTIIs supported a hypothesis in which CXCL13-expressing FDCs in follicles competed with CXCL9/10-expressing monocytes to influence CD4^+^ T cells, the roles of CXCR3 and CXCR5 had not been directly tested [7]. Spatial locations of multiple immune cells and cell-states cannot be easily tested via conventional microscopy due to challenges in imaging tens of surface and intra-nuclear proteins marking heterogeneous CD4^+^ T cell subsets, B cells, monocytes, and other potentially interacting cell-types. Therefore, in this study we tested the capabilities of *Slide-seqV2*, in combination with droplet-based scRNA-seq and computational modelling, to define the phenotypes, positions and co-localisation of numerous transcriptomic states of CD4^+^ T cells, B cells, myeloid cells, and stromal cells in the spleen prior to and during malaria.

## Results

### Slide-seqV2 reveals splenic microarchitecture before and during malaria infection

To systematically examine the phenotypes, locations and possible cellular interactions of CD4^+^ T cells and other immune cells in the spleen during experimental malaria, we first tested *Slide-seqV2*, our spatial transcriptomic technique of 10μm resolution [7, 34], for its capacity to reveal splenic microarchitecture. *Slide-seqV2* comprises a fixed circular array (3mm diameter), or “puck”, of closely packed, positionally-barcoded beads, each 10μm in diameter. Beads capture mRNA locally, allowing for whole transcriptome assessment of tissue at near single-cell resolution. We applied *Slide-seqV2* to mouse spleens prior to and 7 days after *Pc*AS infection (2 pucks per spleen for 2 naïve and 2 infected mice; Fig. 1A), with parallel droplet-based scRNA-seq datasets also generated from the same samples. Flow cytometric assessment confirmed expected levels of parasitemia in the blood (Extended Data Fig. 1A). Analysis of polyclonal CD4^+^ T cells and PbTII cells (Extended Data Fig. 1B) confirmed expected upregulation of CXCR5 and CXCR6 (consistent with Th1/Tfh differentiation having occurred) (Extended Data Fig. 1C-D). In addition, flow cytometric analysis of B cells (Extended Data Fig. 1E) confirmed plasmablast differentiation (Extended Data Fig.1F), which guided selection of samples subjected to *Slide-seqV2*.

**Figure 1.**
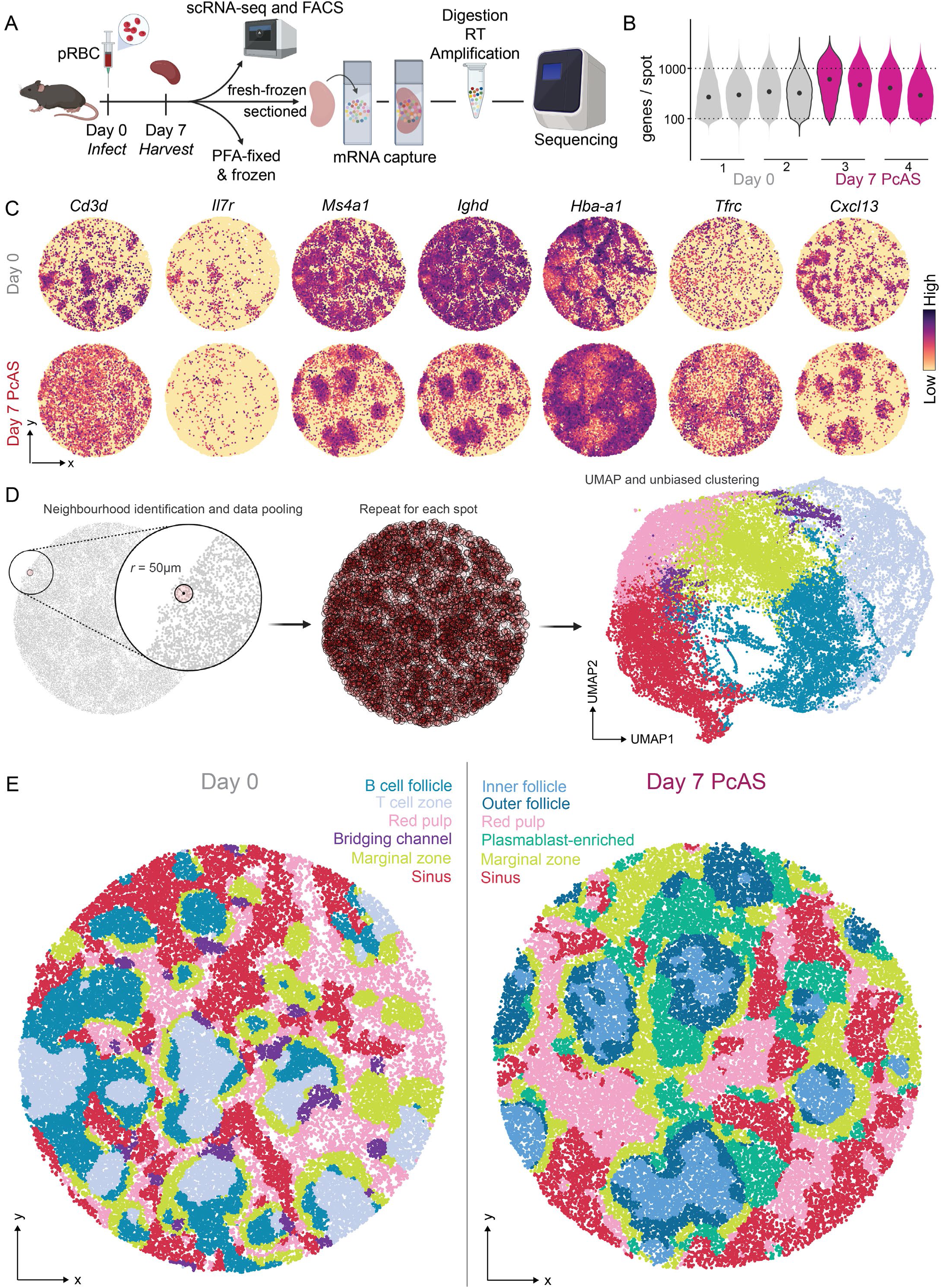
Slide-seqV2 reveals splenic microarchitecture before and during malaria infection. (A) Schematic of *Slide-seqV2* data generation. Uninfected (n=2) or *Pc*AS-infected (n=2) spleens at day 7 post-infection were selected and used for FACS validation, scRNA-seq reference datasets, PFA fixed for banking, or fresh-frozen for spatial transcriptomic analysis using *Slide-seqV2.* Created with biorender. (B) Violin plot of mouse genes detected per spot, per array, per mouse, per timepoint. Highlighted violin plots (with black border) show the arrays selected for display (puck 4 from day 0, puck 1 from day 7). (C) Normalised expression of genes indicating T cells (*Cd3d*, *Il7r*), B cells (*Ms4a1, Ighd*), red blood cells (*Hba-a1, Tfrc*) and stromal cells (*Cxcl13*) in representative arrays in naïve and *Pc*AS-infected spleens. (D) Schematic explaining spatial pooling technique to achieve local neighbourhood-averaging (LNA), unbiased clustering, and UMAP. Beads are contextualised in spatial neighbourhoods of 50μm radii, from which PCA data are pooled inwards and averaged. The resulting data is used as input for UMAP and unbiased clustering (shown). Created with Biorender.com. (E) Unbiased clustering based on technique in (D) for representative Day 0 (left) and Day 7 (right) pucks. Displayed clusters are curated and annotated based on gene expression from raw clustering output in (Extended Data Fig. 3B).

Initial quality control of mouse mRNA from *Slide-seqV2* pucks revealed 68.7% of beads across naïve spleen pucks and 78.0% of those from infected spleen pucks captured at least 100 unique mRNA molecules (Unique Molecular Identifiers, UMIs). We removed from subsequent analysis any beads below this threshold. Post-filtering, a median of 311 (maximum: 7017) genes/bead were detected across naïve pucks, with a median of 424 (maximum: 4385) genes/bead across infected pucks (Fig. 1B, Extended Data Fig. 2A). Several genes commonly expressed by T cells (*Cd3d, IL7r*), B cells (*Ms4a1, Ighd*), red blood cells (*Hba-a1*, *Tfrc*) or stromal cells *(Cxcl13)* appeared to reveal T cell zones, B cell follicles, and red pulp across all pucks, suggesting that *Slide-seqV2* data alone had captured splenic structures (Fig. 1C, Extended Data Fig. 2B). In addition, mapping reads to the *Pc*AS genome revealed small numbers of parasite genes (81.2% of beads from day 7 pucks had at least one *Pc*AS gene detected), particularly in those areas enriched for red blood cells (Extended Data Fig. 2C-D), suggesting *Slide-seqV2* could be employed for studying host-parasite transcriptional interactions in tissue.

Using *Slide-seqV2* data, we next assessed splenic architecture. Unsupervised clustering of beads based on Principal Components, as in standard scRNA-seq analysis pipelines such as *Seurat* [35], provided a coarse-grained depiction of microarchitecture (Extended Data Fig. 3A). However, given that microarchitectural structures are composed of heterogeneous cell types, we reasoned that incorporating gene expression at single spots with those of surrounding spots, via an analysis we termed Local Neighbourhood Averaging (LNA), inspired by *stLearn* [36], would generate spatially-informed mapping (Fig. 1D). This allowed us to group spots with disparate transcriptomes that nevertheless occupied transcriptomically similar niches, which in uninfected spleens resolved expected splenic structures such as T cell zones, B cell zones, marginal zones, and red pulp (Fig. 1E, Extended Data Fig. 3B). At day 7 post-*Pc*AS infection, infection-induced microarchitecture was evident, including differentiated inner and outer sections of B cell follicles, and plasmablast-enriched regions within the red-pulp (Fig. 1E, Extended Data Fig. 3B). These data revealed that *Slide-seqV2* combined with LNA had described expected changes in micro-architecture in a densely packed secondary lymphoid organ.

### Integrating scRNA-seq and *Slide-seqV2* reveals locations of splenic immune and stromal cells

The fixed geometry of beads comprising *Slide-seqV2* pucks resulted, as expected [31, 32], in the capture of mRNAs from different cells on the same bead (Extended Data Fig. 4A). To determine the individual cell types contributing mRNA to any given spot and the proportion of that contribution, *Slide-seqV2* data was integrated with “ground-truth” scRNA-seq reference data, a process termed “*cell-type mapping*” or “*deconvolution*” [37, 38], using *Robust Cell Type Decomposition* (RCTD) (Fig. 2A) [37, 38]. Here, we integrated *Slide-seqV2* data with a whole splenocyte scRNA-seq reference generated from the same tissue (Fig. 2B), and our previous dataset of stromal cells from naïve mouse spleen [39] (Fig. 2C). This facilitated mapping of subsets of B and T cells, as well as B cell zone stroma, termed follicular dendritic cells (FDCs); T cell zone fibroblastic reticular cells (FRCs); red pulp reticular cells (RPRCs); and adventitial reticular cells (ARCs) [39]. To assess the quality of mapping/deconvolution, we defined circular neighbourhoods, 50μm in radius, around each bead and determined which cell types correlated in abundance across those neighbourhoods (Fig. 2D) - 50μm radii were selected to minimise the chance of neighbourhoods spanning different splenic compartments. This revealed expected spatial correlations among cells in shared locations such as B cell subsets and FDCs in B cell zones; T cell subsets and FRCs in T cell zones; and several myeloid cell subsets, erythrocytes, and RPRCs in red pulp (Fig. 2D-F, and Extended Data Fig. 4B). We also observed cell types with ambiguous or in some cases bimodal spatial distributions, including conventional type 1 dendritic cells (cDC1s), plasmacytoid dendritic cells (pDCs), Antibody-Secreting B cells (ASCs), and ARCs (Fig. 2D). Importantly, naïve CD4^+^ T cells, B cells, and monocytes localised to distinct regions, according to existing dogma, which also corresponded to the regions identified via unbiased clustering (Fig. 1E). Thus, integration of scRNA-seq reference data and *Slide-seqV2* data enabled fine mapping of immune and stromal cell populations into expected regions within the steady-state, naïve mouse spleen.

**Figure 2.**
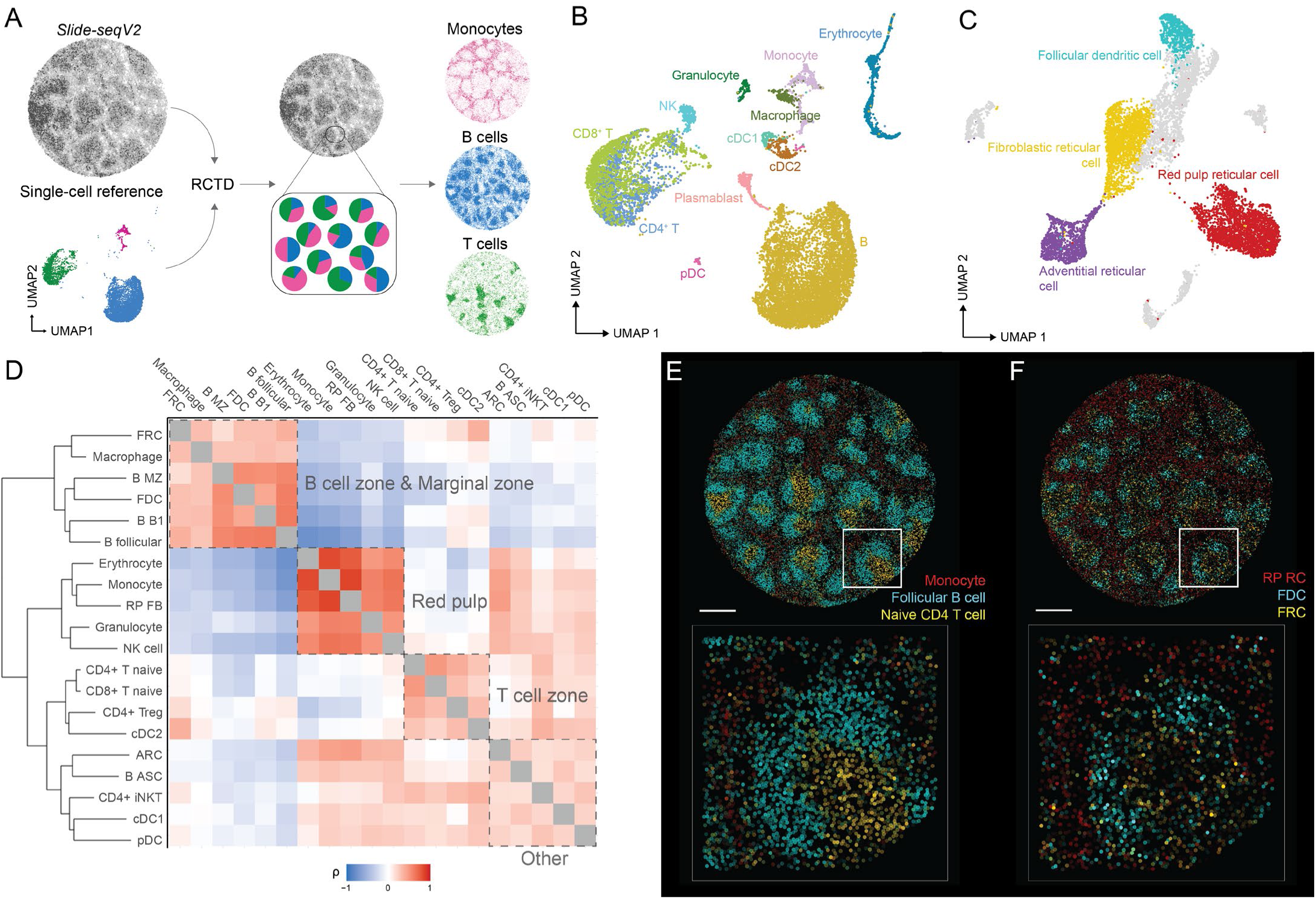
Integrating scRNA-seq and Slide-seqV2 reveals locations of splenic immune and stromal cells. (A) Schematic of single cell and spatial transcriptomic data integration via Robust Cell Type Decomposition (RCTD) to define spatially confined microanatomical regions in the spleen. (B) UMAP of immune cell types identified by high-dimensional unsupervised clustering analysis among day 0 and day 7 splenocyte scRNA-seq transcriptomes. (C) UMAP of stromal cell types used as a reference for deconvolution, curated from Alexandre et al. [48]. (D) Spatial correlations among all day 0 splenic cell-types calculated from randomly selected 50μm radius neighbourhoods. Cell types ordered by hierarchical clustering. Anatomical regions annotated manually. Dendrogram shown at left. Colours reflect Spearman’s rho. (E) RCTD-inferred locations of naïve CD4^+^ T cells (gold), follicular B cells (cyan), and monocytes (red) in a whole day 0 puck with inset showing a representative follicle and T cell zone, indicated by white border above. Scale bar shown is 500µm.

### Mapping B cell differentiation reveals a prolonged bystander response in splenic follicular B cells

To examine the spatial locations of multiple immune cell-types and cell states during infection, it was necessary to deconvolute *Slide-seqV2* pucks with ‘ground-truth’ scRNA-seq reference data from infected spleens. Since our whole spleen scRNA-seq reference contained only 100-200 transcriptomes for those cell-states of most importance for this study, particularly of activated CD4^+^ T cell and B cell subsets (Fig. 2B), we required larger and more representative B cell and polyclonal CD4^+^ T cell scRNA-seq reference datasets. We first focused on B cells because there were no publicly available scRNA-seq datasets for splenic B cells during experimental malaria. In addition, while annotating our initial whole spleen scRNA-seq reference we noted unexpected possible transcriptomic change in all follicular B cells that nevertheless lacked other markers of activation (Extended Data Fig. 5A-B). We hypothesized possible technical batch effects in our data and sought to test this in generating a new comprehensive temporal map of B cell differentiation over the first two weeks of infection. Splenic CD19^+^ B220^hi-int^ cells were sorted prior to and at days 4, 7, 10, and 14 post-infection and processed via droplet-based scRNA-seq (Fig. 3A, and Extended Data Fig. 5C).

**Figure 3.**
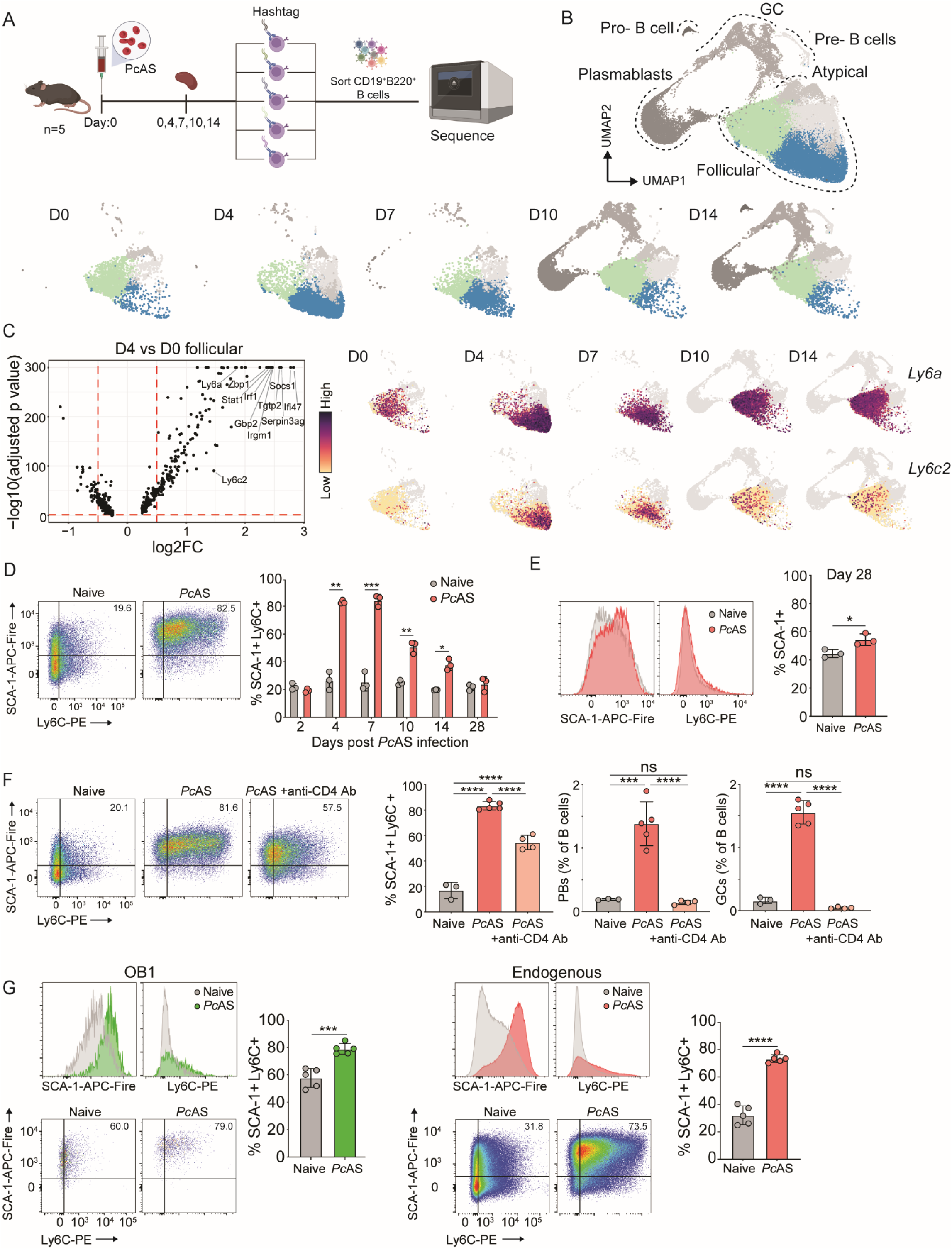
Mapping B cell differentiation reveals a prolonged bystander response in splenic follicular B cells. (A) Schematic of scRNAseq experiment to assess the follicular B cell dynamics over the course of a *Pc*AS infection. C57/BL6 mice were infected with *Pc*AS parasitized red blood cells and splenic B cells were sorted from 5x C57/BL6 mice per timepoint post-infection. Each replicate mouse per timepoint were barcoded with unique TotalSeq™–C hashtags prior to loading samples into 10X Chromium Controller. Created with Biorender.com. (B) High dimensional clustering showing follicular (coloured) and immature and mature-like B cells (grey) represented by the combined UMAP and UMAPs split by timepoints (D0-D14). (C) Volcano plot highlighting the top differentially upregulated genes expressed in the D4 (dark blue) follicular cluster compared to D0 (light blue) follicular cluster plus feature plots of cell surface markers *Ly6a* and *Ly6c2*. (D) Representative FACS plot of SCA-1 and Ly6C expression in IgD^+^ B cells at D4 post *Pc*AS infection and bar graph of the frequency of SCA-1^+^ Ly6C^+^ cells amongst IgD^+^ B cells over the course of a *Pc*AS infection. Statistical test used is multiple Welch’s t-tests, n=3 biological replicates. Data representative of two independent experiments. (E) Representative FACS histograms of SCA-1 and Ly6C expression and bar graph of proportion of SCA-1^+^ in day 28 post-infected IgD^+^ B cells. Statistical test used is Welch’s t-test, n=3 biological replicates. Data representative of two independent experiments. (F) Representative FACS plots and bar graph of SCA-1 and Ly6C expression on IgD^+^ B cells as well as frequency of plasmablast and germinal centre B cells relative to all B cells 7 days post *Pc*AS infection, with or without CD4 T cell depletion. Statistical test used is one-way ANOVA with Tukey’s multiple comparisons test, n=3-5 biological replicates. Data representative of two independent experiments. (G) Representative FACS histograms and pseudocolor plots bar graphs showing frequency of SCA-1 and Ly6C expression on CD86^lo^ IgD^+^ endogenous and CD86^lo^ OB1 transferred B cells, 4 days post *Pc*AS infection. Statistical test used is Welch’s t-test, n=5 biological replicates. Data representative of two independent experiments. Data are ± S.D. p-value * <0.05,

After quality control for high quality B cell transcriptomes (Extended Data Fig. 5D), and hashtag deconvolution of individual mouse samples (Extended Data Fig. 5E), 47,364 cells were advanced for further analysis. Principal Component Analysis (PCA) followed by Uniform Manifold Approximation Projection (UMAP) and unsupervised clustering revealed the emergence over time of a cluster of B-cells expressing high levels of *Xbp1, Sdc1* (encoding CD138), and *Ighg2c*, consistent with being class-switched plasmablasts (Extended Data Fig. 5F). These cells peaked by day 10 p.i., consistent with flow cytometric assessment (Extended Data Fig. 5G). In addition, from day 10 and increasing by Day 14, we noted clusters expressing *Aicda* and *S1pr2*, consistent with annotation as GC B cells (Extended Data Fig. 5F). Finally, a small cluster of cells co-expressed *Tbx21, Itgax* and *Cxcr3*, consistent with being atypical B cells (Extended Data Fig. 5F). Hence scRNA-seq analysis had detected transcriptomes for activated B cell types commonly known to differentiate during experimental malaria. In addition, as suggested in our whole spleen reference data (Extended Data Fig. 5A), most follicular B cells (expressing *Fcer2a* and not isotype-switched (Extended Data Fig. 5F)) shifted from one main transcriptomic cluster prior to infection, to another cluster by days 4 and 7 post-infection, with partial reversion by day 14 p.i. (Fig. 3B). Differential gene expression analysis at day 4 p.i. *versus* follicular cells prior to infection (day 0) revealed up-regulation of genes including *Socs1, Irf1* and *Irgm1* (Fig. 3C), and those enriched for interferon response pathways (Extended Data Fig. 5H). Among upregulated genes, *Ly6a* (encoding SCA-1) remained elevated until day 14, while *Ly6c2*, encoding Ly6C, returned to baseline levels (Fig. 3C). These dynamics were validated at protein level for SCA-1 and Ly6C, with co-expression by a large proportion of splenic IgD^+^ B cells by day 4 of infection, which was sustained until day 14 (Fig. 3D), with SCA-1 remaining elevated at day 28 post infection (Fig. 3E). Together, these data revealed activation of most naïve follicular B cells during *Pc*AS infection marked by SCA-1/Ly6C upregulation. Notably, SCA-1/Ly6C up-regulation on all follicular B cells still occurred in CD4-depleted mice (Fig. 3F and Extended Data Fig. 5I), while plasmablast differentiation was abrogated (Fig. 3F), as previously reported [14]. To determine if SCA-1/Ly6C upregulation occurred in B cells with no specificity to *Plasmodium* antigens, we adoptively transferred ovalbumin-specific OB1 cells prior to *Pc*AS infection. By 4 days p.i. OB1 cells also substantially upregulated SCA-1 and Ly6C compared to uninfected controls, similar to endogenous B cells (Fig. 3G). Thus, scRNA-seq not only generated a temporal map of B cell differentiation during malaria, but also revealed a prolonged, largely CD4^+^ T cell independent, antigen independent bystander B cell response during *Pc*AS infection marked by SCA-1 and Ly6C upregulation.

### Single polyclonal CD4^+^ T cells differentiate primarily into Th1 and Tfh during experimental malaria

We next required a scRNA-seq reference dataset for polyclonal CD4^+^ T cell transcriptomes during malaria, since the initial whole spleen reference contained few Th1 (∼100 cells) or Tfh-like (∼175 cells) polyclonal CD4^+^ T cell transcriptomes. Moreover, our previous studies had focussed solely on TCR transgenic PbTII cells [7, 34], raising the question of their relevance to TCR diverse responses [40]. Therefore, we sought to both generate a polyclonal CD4^+^ T cell scRNA-seq reference, and to determine how the breadth of these compared to PbTII cells.

To examine *Plasmodium*-specific polyclonal CD4^+^ T cells without narrowing the breadth of antigen-specificities studied (*e.g.* via MHC-II tetramers), we opted to sort CD11a^hi^CXCR3^+^ cells (Fig. 4A), since these appeared to contain all Th1 (CXCR6^+^Tbet^+^) and Tfh-like (CXCR5^+^Bcl6^+^) cells (Fig. 4B), as well as capturing higher frequencies of CXCR5^+^ cells compared to a previous approach of examining CD11a^hi^CD49d^+^ cells (Extended Data Fig. 6A-B) [41]. Splenic CD11a^hi^CXCR3^+^ CD4^+^ polyclonal T cells and GFP^+^ PbTII cells were recovered 7 days p.i. (Extended Data Fig. 6C), pooled and processed via droplet-based scRNA-seq (Fig. 4C). After quality control for high-quality transcriptomes and deconvolution of individual mouse samples (Extended Data Fig. 7A-B), 11,529 polyclonal and 1,183 PbTII cells were selected for downstream analysis. After eGFP, TRA and TRB gene removal to focus on transcriptomic heterogeneity (Extended Data Fig. 7C), and removal of antigen presenting cells (Extended Data Fig. 7D) and NKT cells (Extended Data Fig. 7E), PCA followed by UMAP based clustering suggested two main transcriptomic regions occupied by polyclonal cells, with PbTII transcriptomes directly overlapping with these, but importantly not all sub-regions (Fig. 4D). PbTII cells overlapped with polyclonal cells in the areas dominated by *Mki67, Cxcr6/Tbet* or *Cxcr5/Bcl6* co-expression, suggesting that previously reported transcriptomes of proliferating and Th1/Tfh-like PbTIIs were well-represented amongst polyclonal counterparts (Fig. 4D and E).

**Figure 4.**
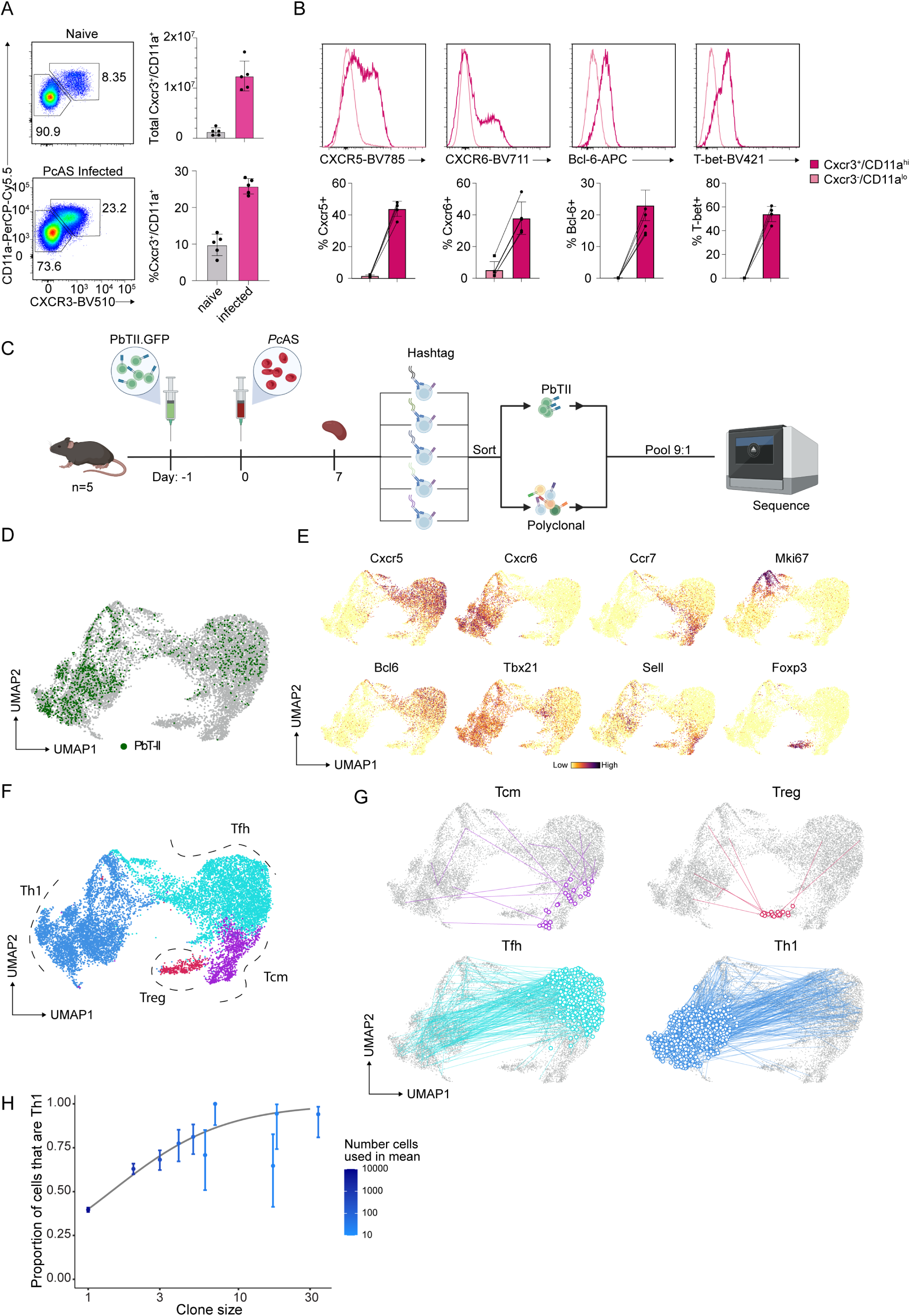
Single polyclonal CD4+ T cells differentiate primarily into Th1 and Tfh during experimental malaria. A) Representative FACS plots of CXCR3 and CD11a expression by CD4^+^ T cells in naïve and infected mice seven days p.i. with *PcAS*, and bar graphs showing frequencies of CD11a^hi^CXCR3^+^ cells amongst total CD4^+^ T cells in naïve and infected mice. Data is representative of at least three independent experiments (5 mice per group, per experiment). (B) Representative histograms and bar graphs showing frequencies of CXCR5^+^, CXCR6^+^, Bcl6^+^ and Tbet^+^ cells in CD11a^hi^CXCR3^−^ and CD11a^hi^CXCR3^+^ CD4^+^ T cells. Data is representative of at least three independent experiments (5 mice per group, per experiment). (C) Schematic representation of the experiment. Five C57BL/6 mice received adoptive transfer of GFP^+^PbTII cells one day prior to infection with *PcAS* parasitized red blood cells with spleens harvested seven days after infection. Samples from each mouse were barcoded with unique TotalSeq™-C hashtags. CD11a^hi^CXCR3^+^ and GFP^+^PbTII cells were then sorted and pooled at a proportion of 9:1, respectively, prior to loading the samples into the 10X Chromium Controller. Created with Biorender.com. (D) UMAP visualisation of PbTII and CD11a^hi^CXCR3^+^ cell transcriptomes on day 7 after *PcAS* infection. € UMAP visualisation showing *Cxcr5, Cxcr6, Ccr7, Mki67, Bcl6, Tbx21, Sell* and *Foxp3* on all CD11a^hi^CXCR3^+^ cells on day 7 p.i. (F) UMAP visualization of all CD11a^hi^CXCR3^+^ cells coloured according to cell subtypes; T cell memory precursors (Tcmp), regulatory T cells (iTreg), Follicular helper cells (Tfh) and T helper-1 cells (Th1). (G) UMAP visualisation of polyclonal CD11a^hi^CXCR3^+^ cells with connecting lines linking cells which share identical T cell receptors (TCR). For each UMAP, only clonotypes which contain a cell classified as Tcm-like, Treg, Tfh or Th1 are shown. (H) Linear model of the frequency of Th1 cells *versus* the number of members in the clonotype families identified (clone size). Dots are the proportion of cells that were Th1 of all cells from a given clone size. Error bars indicate the 95% CI (from a binomial distribution) of these proportions. The opacity of each point indicates the number of cel€s identified as being from a clone of that size. The grey line if 36itfitted generalised linear model (with logit link function).

In contrast to the dominant Th1/Tfh phenotypes displayed by polyclonal CD4^+^ T cells and PbTII cells, polyclonal clusters marked by either *Foxp3* expression, or by high levels of *Ccr7/Sell* co-expression, were largely devoid of PbTII cells (Fig. 4D-F), suggesting polyclonal cells could have acquired states not evident in PbTIIs - these clusters exhibited high Treg and Tcm-like scores, respectively, when tested using reported signatures [42, 43] (Extended Data Fig. 8A and B). We then confirmed at protein level that CCR7^+^ CD62L^hi^ Tcm-like cells and CD25^+^ Foxp3^+^ CD4^+^ T cells were detected amongst polyclonal CD11a^hi^CXCR3^+^ CD4^+^T cells but not amongst PbTII cells (Extended Data Fig. 7F and G). Thus, polyclonal CD4^+^ T cells appeared to have differentiated mostly into Th1 or Tfh cells, and possibly, though more rarely, into iTreg or Tcm-like states during infection, while PbTII cells had been similarly confined to Th1/Tfh bifurcation as previously reported [7]. Interestingly, there appeared a transcriptomic continuum between Tfh-like and Tcm-like states, suggesting a complex relationship between the two amongst polyclonal CD4^+^ T cells. Parallel analysis of TCR sequences revealed instances, albeit rare compared to Th1 and Tfh clonal sharing, where Tcm-like and iTreg-like polyclonal cells shared TCR sequences with either Th1 or Tfh cells (Fig. 4G). This suggested that naïve polyclonal CD4^+^ T cells gave rise predominantly to both Th1 and Tfh fates during *Pc*AS infection, and rarely to Tcm-like and iTreg states.

Finally, given that TCR sequence can influence effector fate, we determined whether there was a propensity for individual clones to be dominated by one state over another, a situation we had not observed for PbTII cells [34]. Fate choices of Tcm-like and iTreg states were too infrequent in our datasets to facilitate modelling. Hence, we confined our analysis to Th1 versus Tfh fate bifurcation. Of all cells where TCR sequences had been detected, 3518 of 7806 (45%) of cells were Th1, revealing a relatively equal split between Th1 and Tfh fates at a population level. However, amongst clonally-expanded cells (*ie.* those sharing a TCR with at least one other cell), the frequency of Th1 fate rose to 67.5%. Using a generalised linear model, we observed a significant correlation between the size of a clonal family and the proportion of cells within that family exhibiting a Th1 over a Tfh fate, with a 2.2-fold increase (95% CI: 2.0-2.4) in the odds of a cell being Th1 for every 2-fold increase in family size (p<0.0001) (Fig. 4H). Thus, although it was most common for single naïve CD4^+^ T cells to give rise to both Th1 and Tfh states during experimental malaria, as previously reported for other infections, the most expanded clones were heavily skewed towards a Th1 fate (Fig. 4H). Thus, during experimental malaria in mice, most splenic naïve polyclonal CD4^+^ T cells give rise to both Th1 and Tfh fates, with Th1-bias being evident amongst the most expanded clones.

### Spatial transcriptomic co-localisation analysis predicts cellular and molecular interaction partners for Th1 and Tfh cells

Having assembled scRNA-seq references for immune cells in the spleen, including specific references for polyclonal CD4^+^ T cells and B cells, we next sought to define polyclonal Th1 and Tfh locations and cellular interactions in the spleen during *Pc*AS infection. We hypothesized that polyclonal Th1 cells were attracted to and supported by monocytes found in the red pulp, since monocyte depletion had previously led to reduced Th1 differentiation in PbTII cells [7]. Conversely, we hypothesized that polyclonal Tfh-like cells were located in B cell follicles, prior to the development of germinal centres (GCs). To test these, we employed *RCTD* to deconvolute *Slide-seqV2* pucks at day 7 p.i. (Fig. 5A, and Extended Data Fig. 9). We assessed *RCTD* output by calculating spatial correlations of all cell-types with one another in circular neighbourhoods of 50μm radii. Firstly, relative to naïve mice (Fig. 2D), splenic microarchitecture exhibited an expected reduced distinction between T cell zones and B cell zones, and movement of various lymphocyte subsets to the red pulp (Fig. 5A). Importantly, consistent with our hypotheses, Th1 cells appeared co-located with monocytes in the red pulp, while polyclonal Tfh-like cells co-located with follicular B cells and activated B cells (Fig. 5A-C). In addition, Tcm-like states adopted a similar localisation to Tfh cells (Fig. 5D). These data indicated firstly that polyclonal CD4^+^ T cells exhibiting Th1-like transcriptomes are indeed located adjacent to monocytes in the red pulp, consistent with direct interaction being required for promoting or sustaining a Th1-phenotype. Secondly, our analysis confirms the locations of polyclonal Tfh-like and Tcm-like cells within or close to B cell follicles prior to emergence of GCs. Importantly, our findings relating to splenic microarchitecture and Th1/Tfh localisation using *RCTD* were largely repeated using a second computational approach, *cell2location* (Extended Data Fig. 10A-B) [37, 38]. Finally, our spatial transcriptomic screen predicted Treg cells relocate from T cell zones to the red pulp during infection and localise closely with plasmablasts (Extended Data Fig. 11A). Given their proximity to plasmablasts, we hypothesised that Tregs could influence these cells and performed an unbiased, transcriptome-based, ligand-receptor interaction screen with *CellChat* [44]. We inferred several interactions, such as via *Il10* to *Il10r* and *Entpd1* to *Adora2a* (Extended Data Fig. 11B), which highlights the value of this approach for hypothesis generation and in searching for novel cellular interactions.

**Figure 5.**
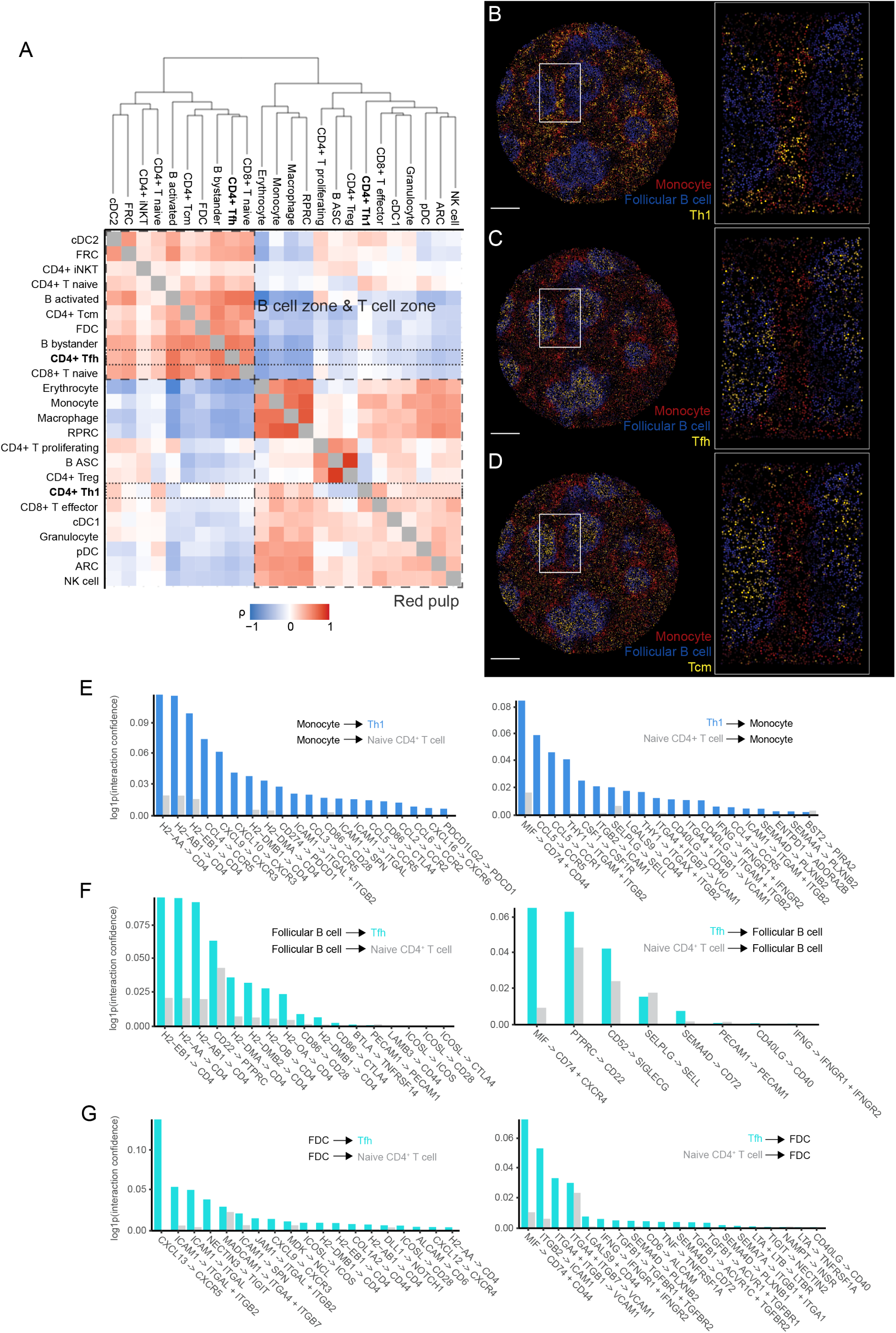
Spatial transcriptomic co-localisation analysis predicts cellular and molecular interaction partners for Th1 and Tfh cells. (A) Spatial correlations among all day 7 splenic cell types calculated from randomly selected 50μm radius neighbourhoods. Cell types ordered by hierarchical clustering. Clusters labelled as B cell and T cell zone or red pulp. Dashed lines indicate CD4+ Tfh and Th1 cells. (B-D) RCTD-inferred locations of Th1 (B), Tfh (C), and Tcmp (D) (gold); follicular B cells (blue); and monocytes (red) in a whole day 7 puck with inset showing representative follicles and interfollicular regions. Colour scale for T cell signatures condensed for visualisation purposes. Scale bar shown is 500€ (E) Ligand-receptor interactions from monocytes to Th1 (left) and Th1 to monocytes (right) ranked by confidence (blue). Confidence values for same ligand-receptor interactions are shown between naïve CD4^+^ T cells and monocytes (grey). (F) Ligand-receptor interactions from follicular B cells to Tfh (left) and Tfh to follicular B cells (right) ranked by confidence (cyan). Confidence values for same ligand-receptor interactions are shown between naïve CD4^+^ T cells and follicular B cells (grey). (G) Ligand-receptor interactions from FDCs to Tfh (left) and Tfh to FDCs (right) ranked by confidence (cyan). Confidence values for same ligand-receptor interactions are shown between naïve CD4^+^ T cells and FDCs (grey).

To determine further receptor/ligand interactions between pairs of co-localised cell-types, namely Th1/monocytes, Tfh/FDC, and Tfh/follicular B cells, we conducted another unbiased screen of cell-cell interactions using *CellChat* [44]. In infected spleens, ligands signalling from monocytes to Th1 cells included MHC-II and chemokine ligands *Ccl4, Cxcl9*, and *Cxcl10*, while those produced by Th1 cells for interaction with monocytes included *Cd40lg, Ifng, Ccl5*, and *Mif* (signalling via *Cd44*) (Fig. 5E) - lower levels of signaling was inferred between monocytes and naïve CD4^+^ T cells. Therefore, spatial data predicted the emergence of antigen-presenting-cell-like interactions via MHCII/TCR and co-stimulatory molecules between Th1 cells and monocytes, as well as chemokine signalling via *Ccl5/Ccr5* and *Cxcl9/Cxcl10/Cxcr3*. Predicted interactions between Tfh and B cells included *Cd40/Cd40lg*, *Mif*/*Cxcr4*, *Icos*/*Icosl*, and MHCII interactions, some of which were also inferred between naïve CD4^+^ T cells and B cells with lower confidence (Fig. 5F). Finally, between FDC and Tfh cells, *CellChat* predicted chemokine signalling via *Cxcr5/Cxcl13* and *Cxcr4/Cxcl12*, and several cell adhesion molecules including *Icam1, Col1a2, Mcam1*, and *Nectin3* (Fig. 5G), with *Cxcl13*-mediated signalling absent between FDCs and naïve CD4^+^ T cells (Fig. 5G). Thus, our spatial inference of molecular interactions was consistent with chemokine-signalling by monocytes and B cell stroma influencing the biology of CD4^+^ T cell differentiation in the spleen, alongside conventional MHCII and co-stimulatory signals.

Taken together, untargeted co-localisation analysis of multiple cell-types and cell states provided evidence consistent with our hypotheses that monocytes support polyclonal Th1 differentiation via MHCII interaction [7], CD40-mediated co-stimulation and CXCR3 and/or CCR5 chemokine signalling, while Tfh cells interact with both B cells and associated B cell stroma via similar though in some cases distinct pathways.

### CXCR5 promotes CD4^+^ T cell expansion during experimental malaria

Given colocalization of Th1 cells with monocytes in the red pulp, positioning of Tfh-like cells within B cell follicles, and predicted roles for chemokine signalling between these cell-types, we next tested our previous hypothesis [7], that primed CD4^+^ T cells experienced competing chemoattraction from CXCL13^+^ FDCs versus CXCL9/10-expressing monocytes due to early co-expression of CXCR3 and CXCR5. To test this *in vivo*, naïve PbTII cells were genome-edited via CRISPR/Cas9 with guide RNAs targeting either or both *Cxcr3, Cxcr5* (or control *Cd19*), and assessed for clonal expansion and Th1 differentiation at 7 days after *Pc*AS infection (Fig. 6A). Firstly, FACS analysis confirmed that upregulation of CXCR3 or CXCR5 was largely abrogated by specific guide RNAs (Fig. 6B and C). Most strikingly, CXCR5, but not CXCR3, supported PbTII numbers in the spleen (Fig. 6D), suggesting an unexpected role for CXCR5 in clonal expansion. In examining Th1-differentiation, we noted CXCR5, but not CXCR3, exerted a modest suppressive effect on direct *ex vivo* production of IFNγ by PbTIIs (Fig. 6E) compared to endogenous CD4+ T cells (Extended Data Fig. 12A), with evidence of Tbet suppression only when compared to endogenous naïve CD4^+^ T cells (Extended Data Fig. 12B), but no evidence of suppression of another Th1-marker, CCL5 (Extended Data Fig. 12C). These data suggested that although CXCR5 played a modest role in suppressing Th1 differentiation, CXCR3 played no such opposing role. Moreover, we revealed an unexpected role for CXCR5 in supporting clonal expansion of CD4^+^ T cells in the spleen during malaria.

**Figure 6.**
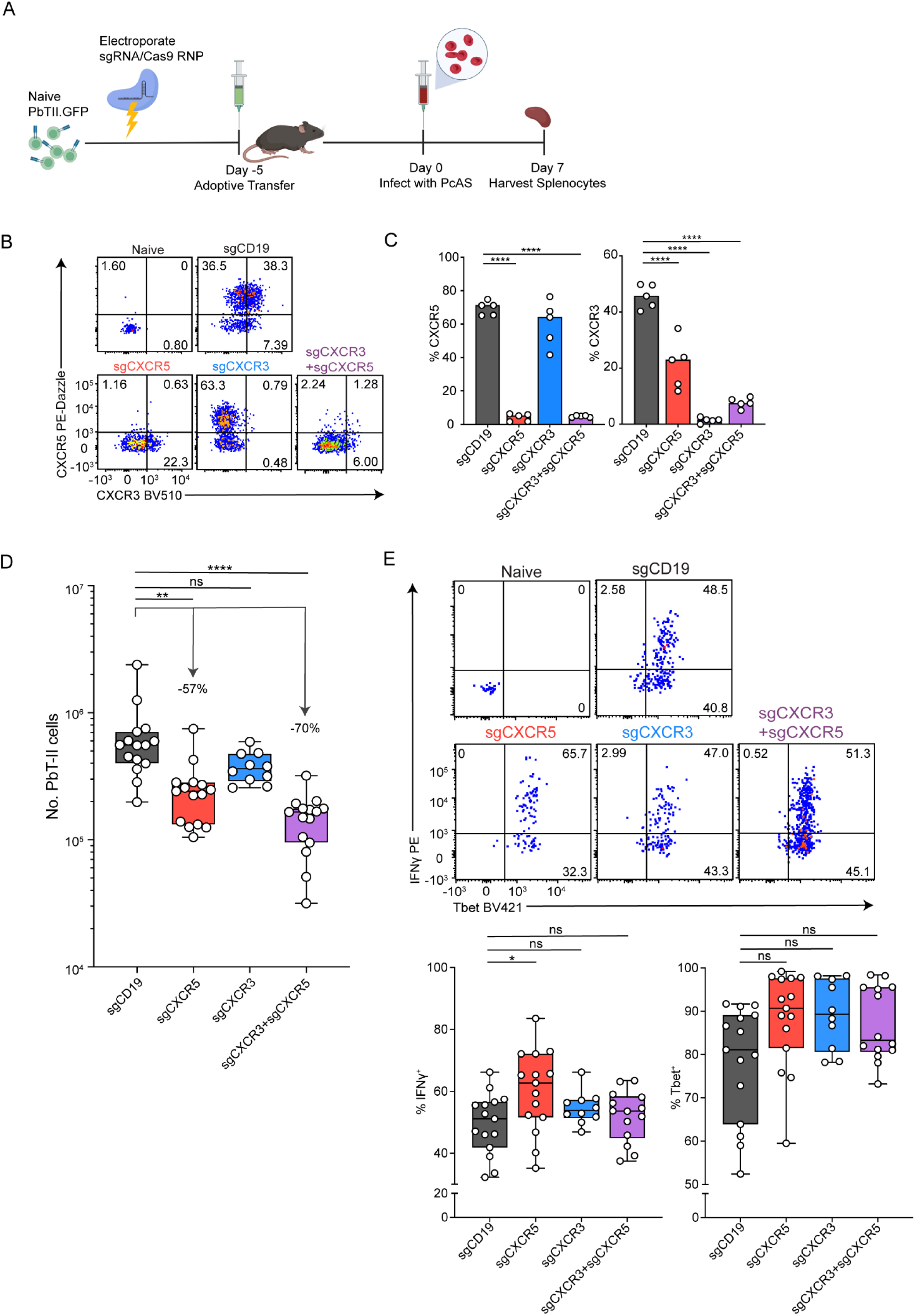
CXCR5 promotes CD4+ T cell expansion. (A) Naïve GFP^+^ PbTII cells were electroporated with sgRNA/Cas9 RNPs and immediately transferred into C57BL/6 recipient mice (Day -5). On day 0, mice were infected with *Pc*AS and splenocytes were analysed on day 7. Created with Biorender.com. (B, C) Representative FACS plots (B), and bar graphs (C) showing the frequencies of CXCR3^+^ and CXCR5^+^ naïve PbTII cells and CRISPR/Cas9 knockout sgCD19, sgCXCR3, sgCXCR5, or sgCXCR3+sgCXCR5 GFP^+^ PbTII cells at day 7 p.i. Data representative of three independent experiments (n=5 biological replicates per group). Statistical test performed using paired two-way ANOVA with Tukey’s multiple comparison test. p-values are indicated where ****p<0.0001. (D) Bar graphs showing total number of GFP^+^ PbTII cells per spleen in the respective CRISPR edited cells at day 7 p.i. (E) Representative FACS plots and bar graphs of direct *ex vivo* staining of IFN-γ^+^ and Tbet^+^ sgCD19, sgCXCR3, sgCXCR5, or sgCXCR3+sgCXCR5 GFP^+^ PbT-ΙΙ cells at day 7 p.i. (D, E) Data are pooled from 3 independent experiments for sgCD19, sgCXCR5, and sgCXCR3+sgCXCR5 GFP^+^ PbTII cells and pooled from 2 independent experiments for sgCXCR3 GFP^+^ PbTII cells (n= 5 mice per group, per independent experiment). Bars indicate median. Statistical test performed using paired two-way ANOVA with Tukey’s multiple comparison test. P-values are indicated where *p< 0.05, **p< 0.01, ****p<0.0001.

## Discussion

During blood-stage infection with malaria parasites, the spleen harbours diverse communities of cell types in various states of activation, often occupying discrete microanatomical niches. To systematically identify all cell types, their transcriptional states, their locations, and possible cell-cell interactions has been challenging, since few platforms have been capable of assessing single cells at genomic scale within tissues. In this study, we tested how *Slide-seqV2*, our untargeted spatial transcriptomic platform at near-single cell resolution, could examine locations and cellular interactions of polyclonal CD4^+^ T cells and B cells as they differentiated in the spleen during experimental malaria in mice. Our central biological goal was to understand how polyclonal CD4^+^ T cells differentiate into Th1 and Tfh cells in the context of other changes occurring in the spleen during infection.

First, we found that *Slide-seqV2* data alone, when analysed using our Local Neighbourhood Averaging (LNA) approach, could readily identify microanatomical niches during steady state and microanatomical change during infection. It is perhaps counterintuitive that by averaging transcriptomic data within a certain area around a given bead, and repeating this iteratively across the tissue, that a more interpretable tissue map was constructed compared to a simple assessment of each bead without averaging. Nevertheless, the high resolution of *Slide-seqV2* permitted such fine analyses of tissue structure. When combined with either LNA here, or perhaps other computational approaches that incorporate other data types such as microscopy images [36], this may pave a way towards automated tissue histology at genome-scale, with hundreds to thousands of genes detected per bead.

Since *Plasmodium* parasites produce polyadenylated mRNAs, we were able to detect *Plasmodium* genes expressed across *Slide-seqV2* pucks, particularly in red pulp areas of the spleen. Although the total number of parasites genes detected across whole pucks was close to genome-scale, it was clear that only a handful of genes were detected per bead, suggesting in this instance *Slide-seqV2* could not assess specific aspects of parasite biology. However, it is possible for certain questions related to protozoan parasites and their host interactions, for example sequestered malaria parasites in brain tissue [45, 46], or intracellular *Leishmania* parasites within tissue lesions, that untargeted spatial transcriptomics may be beneficial [47].

Combining *Slide-seqV2* data with scRNA-seq reference data afforded several benefits. First, we could determine which cell-types contributed mRNA to any given bead, and in what proportions. This process is termed “cell-mapping” or “deconvolution” and permitted *Slide-seqV2* to approximate single-cell resolution. Second, scRNA-seq references enabled identification and localisation of as many cell types and cell states as were present within the scRNA-seq reference. This distinguished subtly different states of one cell type (*e.g.* Tcm-like versus Tfh versus Th1 amongst CD4^+^ T cells), as well as identifying cells for which unequivocal single-gene/protein markers do not exist. Although the requirement for scRNA-seq reference data poses possible financial and logistical constraints, our use of stromal cell references generated previously for a separate study [48] suggests that future studies can use publicly available scRNA-seq data to perform deconvolution and cell-type identification.

A third benefit of utilising scRNA-seq reference data was the increased detail with which any given cell-type or state could be examined, once located in tissue. For instance, given that polyclonal CD4^+^ T cells were found adjacent to monocytes, we examined their transcriptomes in tandem, using *CellChat* [44], for possible receptor/ligand interactions. This suggested MHCII, costimulatory and various cytokine-based interactions. Perhaps most importantly, by screening tens of different cell-types and cell-states within tissue, novel cell-cell interactions could be predicted. For example, we predict based on our analyses, that Foxp3^+^ regulatory CD4^+^ T cells re-locate from the T-cell zone to the red pulp during the first week of experimental malaria, where they can be found adjacent to plasmablasts. Furthermore, *CellChat* predicts possible interactions between these cell-types via CD39/CD73 and IL-10/IL-10R. CD39 and IL-10 signalling by plasmablasts has been described in other disease models [49, 50], but remains unexplored during parasitic infection. Furthermore, possible roles for Foxp3^+^ regulatory CD4^+^ T cells in regulating plasmablast responses remains largely unexplored in any system. A caveat of using scRNA-seq reference data was that the ability to map cells depended on the quality and annotation of the reference itself. In situations where cell-types were complex, for example myeloid cells, this was challenging, and might be improved by using larger datasets, as we did for CD4^+^ T cells and B cells.

*Slide-seqV2* and scRNA-seq identified a follicular bystander B cell response during malaria characterised by upregulation of SCA-1, Ly6C and type I and II interferon response pathways [51–54], all occurring early, by day 4 post-infection, as parasitemia became patent. Given Type I IFNs are detected early in the spleens [55] and plasma [56] of *Plasmodium*-infected mice, we hypothesise that Type I IFN and/or innate immune signalling via TLRs or other receptors directly induce bystander activation in B cells. A similar response was reported in the respiratory tract after influenza infection in mice [57], mediated by Type I IFN, and characterised by CD69 and CD86 expression. The consequences of bystander activation of all follicular B cells during malaria, and whether this might occur in human infection remains unknown. Given the substantial upregulation of *Socs1*, a known suppressor of cytokine signalling [58], and since another *Socs* protein, *Socs3*, acts in B cells to modulate germinal centre and antibody responses *in vivo* [59], we hypothesise that *Socs1* serves to limit unwanted B cell activation during *Plasmodium* infection.

A central goal of our study was to examine CD4^+^ T cell differentiation within the context of all other cellular processes occurring within the spleen, and so discover cell-types influencing this process. In particular, we sought to understand how our previous work using TCR transgenic PbTII cells [7, 34] related to the biology of TCR diverse polyclonal CD4^+^ T cells. Comparison of the two within the same mice revealed strikingly similar transcriptomic profiles for Th1 and Tfh states adopted by PbTII and polyclonal CD4^+^ T cells, with VDJ sequencing also revealing that as with PbTII cells, the majority of responding naïve TCR diverse clones gave rise to both Th1 and Tfh states. In addition, we noted two possible other phenomena absent from PbTII responses. Firstly, we noted rare instances in which some TCR diverse clones had given rise to Foxp3^+^ or Tcm-like phenotypes in addition to Th1 and/or Tfh states. These data suggest that Tcm-like differentiation does occur amongst polyclonal CD4^+^ T cells during *Plasmodium* infection, as previously reported for LCMV-infection [43], but that PbTII cells do not exhibit this phenotype, consistent with our previous studies [7, 34]. A caveat to our experiments is that Tcm-like cells amongst CD11a^hi^CXCR3^hi^ cells may constitute pre-existing Tcm cells un-related to *Plasmodium* infection. The rare instances of clonal sharing between Tcm-like cells and Th1/Tfh cells will need further investigation, ideally with much higher numbers of cells, to determine if this fate choice is real yet rare during experimental malaria. Moreover, our data re-inforce the concept of a transcriptional continuum between Tfh cells and Tcm-like cells, which raises the question of whether and how fate between these two states is determined within the spleen. Given the apparent similar location of these cell-types with the spleen, we speculate CD4^+^ T-cell intrinsic stochasticity may play a role in Tcm-like *versus* Tfh-like bifurcation. The second phenomenon observed amongst polyclonal Th1/Tfh clones was an apparent skewing towards a Th1 fate as clonal family size increased. We hypothesise that TCRs belonging to the most expanded clones may exhibit unique affinities or interactions with MHCII-peptide complexes that both support clonal expansion and either inhibit Tfh differentiation or support Th1 [60, 61]. It is possible that some fully differentiated Th1 clones exhibit a prolonged burst of clonal expansion [62], thus being over-represented within the small pool of cells assessed by scRNA-seq and making it harder to detect Tfh counterparts. Future work aims to clarify the nature of the TCRs encoded by these Th1-dominated expanded clones.

Our previous cellular depletion studies revealed monocytes and B cells supported Th1/Tfh bifurcation in PbTII cells [7, 63]. Here, we provide evidence that monocytes, and cDC2, are indeed located adjacent to polyclonal Th1 cells, and that monocyte upregulation of MHCII, CXCL9/10 and other molecules could mediate direct interactions with Th1 cells via CXCR3. Confirmation of co-location of polyclonal Tfh-like cells with B cell follicles expressing CXCL13 was consistent with our previous model of monocytes and B cells controlling Th1/Tfh differentiation. However, our CRISPR/Cas9 experiments suggested that CXCR3 played no role, and CXCR5 only a modest role in controlling Th1/Tfh bifurcation, and that other factors, such as early paracrine production and response to locally-made IL-2 may control this process [64]. In addition, it was surprising to us that CXCR5 appeared to control the magnitude of clonal expansion of PbTIIs in our model, whereas CXCR3 had no such effect. Mechanisms by which CXCR5 could mediate this effect remain to be uncovered. We hypothesize CXCR5 might support early trafficking of activated CD4^+^T cells to the edges of the T cell zone, and in doing so might create opportunities for APC interactions [65]. Our studies are in contrast to a vaccination study in which adoptively transferred *Cxcr5^−/−^* CD4^+^ T cells expanded as effectively as wild-type cells [66], as well as an LCMV-infection study in which temporally-restricted CXCR5-deficiency, initiated 4-days post-infection, had no effect on CD4^+^ T cell numbers. We speculate that context-specific factors, and timing of CXCR5-deficiency may explain apparent differences between our studies. Finally, reasons for no loss of Th1 differentiation in the absence of CXCR3 remain unclear. It is possible that although monocytes support Th1-differentiation, that CXCR3 is not needed for this process. Instead, given the multiple inferred cell-cell interactions between Th1 cells and monocytes, we hypothesise that combined signalling by CCL2, CCL3, and CCL5 as well as MHCII and co-stimulation might all serve to facilitate monocyte/Th1 interaction. In summary, we have shown that spatial transcriptomics at near single-cell resolution can provide detailed insight into the microanatomy of the spleen as well as the identities, locations and potential cell-cell interactions of tens of different cell-types and cell-states during an immune response. In studying experimental malaria, we reveal novel biological processes in B cells, predict novel interactions between CD4^+^ T cell subsets and activated B cell states or myeloid cells, and provide further insight into mechanisms controlling Th1/Tfh differentiation in the spleen. We anticipate our general approaches and high-resolution datasets will constitute an important resource for future assessments of secondary lymphoid organs in health and disease.

## Methods

### Mice, adoptive transfer and infection

*Mice* - C57BL/6J mice were obtained from Animal Resources Centre (Perth, Western Australia, Australia) and maintained under specific-pathogen-free conditions within the Biological Research facility of the Doherty institute for Infection and Immunity (Melbourne, Victoria, Australia) with a 12 h day/night light cycle, humidity of 40–70%, and temperature of 19–22 °C, or within the Animal Facility at QIMR Berghofer Medical Research Institute (Brisbane, Queensland, Australia) with a 12 h day/night light cycle, humidity of 55–65%, and temperature of 19–22 °C. PbTII.nzEGFP mice and OB1.*Rag*−/−.uGFP mice were bred in-house in the Biological Research Facility of the Peter Doherty Institute for Infection and Immunity and maintained under the same conditions described above. All mice were female, 6-10 weeks old, all procedures were approved by the University of Melbourne Animal Ethics Committee (1915018) and the QIMR Berghofer Medical Research Institute Animal Ethics Committee (A1503-601M).

*Adoptive transfer* – naïve spleens from PbTII.nzEGFP or OB1.*Rag*-/-.uGFP female donors were harvested and homogenized through cell strainers of 70-µm to create a single-cell suspension. Red blood cells (RBCs) were lysed using Lysing Buffer (BD) and the CD4^+^ PbTII cells were enriched using CD4 (L3T4) microbeads (Miltenyi Biotec). Each mouse received 10^4^ PbTII cells or 15×10^6^ OB1 cells injected into the lateral tail veins.

*Infection* – C57BL/6J passage mice were infected using thawed *Pc*AS infected blood stabilites. *Pc*AS-infected RBCs were collected from passage mice and 10^5^ infected RBCs were injected into each mouse via lateral tail vein.

*Parasitemia assessment* – Parasitemia measurements were carried out as described previously [5]. Briefly, a drop of blood was collected from the tail vein into RPMI containing 5U/ml of heparin. Diluted blood samples were stained at room temperature, for 30 minutes in the dark, with a combination of Hoechst 33342 (10µg/ml; Sigma-Aldrich) and Syto84 (5µM; Life Technologies). Staining was quenched with 10 times initial volume of RPMI and percentage of Hoechst^+^ Syto84^+^ RBCs assessed using a Fortessa cytometer (BD).

*CD4^+^ cell depletion –* At 4 days prior to *Pc*AS infection, mice were administered 0.2mg of anti-CD4 depleting monoclonal antibody (clone GK1.5, BioXCell) or isotype control antibody (clone LTF-2, BioXcell) and a further 0.1mg dose was administered one day prior to infection. Antibodies were administered by intra-peritoneal injection in 200ul of sterile PBS.

### Flow Cytometry

Surface staining of mouse splenocytes was performed sequentially, first with Zombie Yellow diluted in phosphate buffered saline (PBS), followed by Fc Receptor blocking using commercial polyclonal antibodies and finally incubation on ice for 20 minutes with different titrated panels of monoclonal antibodies diluted with PBS + 2% fetal bovine serum (FBS) and 2mM EDTA.

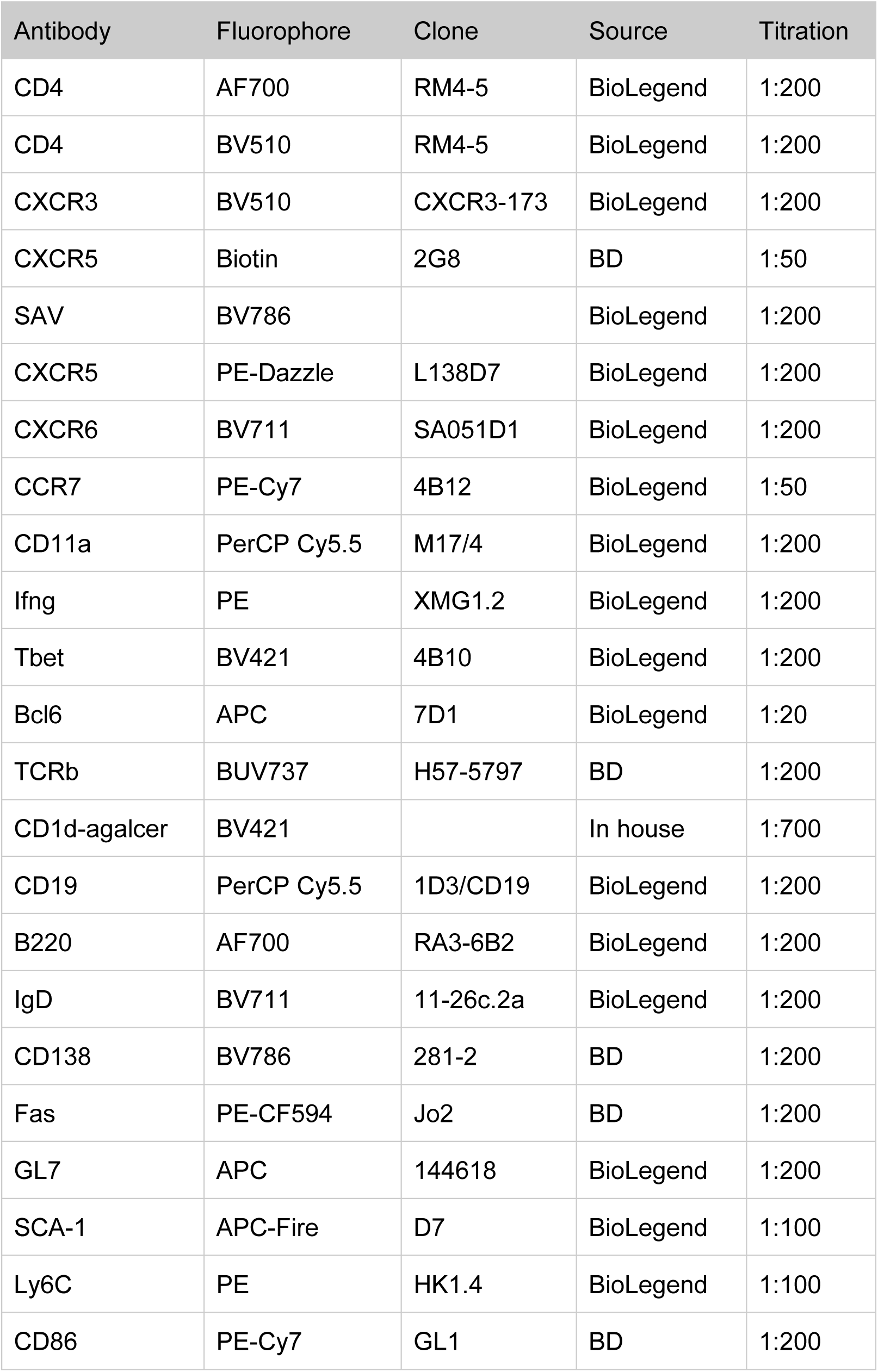

| Antibody | Fluorophore | Clone | Source | Titration |
| --- | --- | --- | --- | --- |
| CD4 | AF700 | RM4-5 | BioLegend | 1:200 |
| CD4 | BV510 | RM4-5 | BioLegend | 1:200 |
| CXCR3 | BV510 | CXCR3-173 | BioLegend | 1:200 |
| CXCR5 | Biotin | 2G8 | BD | 1:50 |
| SAV | BV786 |  | BioLegend | 1:200 |
| CXCR5 | PE-Dazzle | L138D7 | BioLegend | 1:200 |
| CXCR6 | BV711 | SA051D1 | BioLegend | 1:200 |
| CCR7 | PE-Cy7 | 4B12 | BioLegend | 1:50 |
| CD11a | PerCP Cy5.5 | M17/4 | BioLegend | 1:200 |
| Ifng | PE | XMG1.2 | BioLegend | 1:200 |
| Tbet | BV421 | 4B10 | BioLegend | 1:200 |
| Bcl6 | APC | 7D1 | BioLegend | 1:20 |
| TCRb | BUV737 | H57-5797 | BD | 1:200 |
| CD1d-agalcer | BV421 |  | In house | 1:700 |
| CD19 | PerCP Cy5.5 | 1D3/CD19 | BioLegend | 1:200 |
| B220 | AF700 | RA3-6B2 | BioLegend | 1:200 |
| IgD | BV711 | 11-26c.2a | BioLegend | 1:200 |
| CD138 | BV786 | 281-2 | BD | 1:200 |
| Fas | PE-CF594 | Jo2 | BD | 1:200 |
| GL7 | APC | 144618 | BioLegend | 1:200 |
| SCA-1 | APC-Fire | D7 | BioLegend | 1:100 |
| Ly6C | PE | HK1.4 | BioLegend | 1:100 |
| CD86 | PE-Cy7 | GL1 | BD | 1:200 |

For the panels containing CCR7 staining, cells were first incubated with anti-CCR7 antibody for 1 hour, 37°C prior to staining with remaining antibodies. For intracellular staining, surface stained cells were fixed and permeabilised using the eBioscience FoxP3 kit then stained with relevant titrated antibodies for 1 hour on ice. Acquisition of cells was conducted on Fortessa cytometers (BD), with all analysis subsequently done using Flowjo (Treestar). For SCA-1 and Ly6C FACS plots, gates drawn based on FMOs for each marker.

### Preparation of cells for Cell Sorting and scRNA-seq

*Spleen D0 and D7 dataset* - Spleens were harvested and homogenized through 100μm cell strainers to create single-cell suspensions. RBCs were lysed using Lysing Buffer Hybri-Max (Sigma-Aldrich). Samples were washed twice in PBS 2% BSA and resuspended in PBS 2% BSA containing propidium iodide at dilution of 1:500. Live cells were sorted using a BD FACS Aria III and loaded onto the Chromium controller. cDNA sequencing libraries were prepared using Single-cell 3’ reagent kits (10X Genomics).

*T and B cell reference datasets* - Spleens from infected mice and naïve controls were harvested and homogenized through 70µm cell strainers to create a single-cell suspension. Red blood cells were lysed using Lysing Buffer (BD) and the samples were washed twice in phosphate buffered saline (PBS) containing 2% of BSA. Prior to antibody staining, samples were incubated with Fc block reagent (BD). Cells were then incubated with titrated panels of fluorescent monoclonal antibodies and TotalSeq^TM^-C mouse hashtags for 20 minutes on ice.

| Antibody | Fluorophore | Clone | Source | Titration |
| --- | --- | --- | --- | --- |
| CD45/MHC-I | Hashtag_1 | M1/42; 30-F11; | BioLegend | 0.1-0.5µg/million cells |
| CD45/MHC-I | Hashtag_2 | M1/42; 30-F11; | BioLegend | 0.1-0.5µg/million cells |
| CD45/MHC-I | Hashtag_3 | M1/42; 30-F11; | BioLegend | 0.1-0.5µg/million cells |
| CD45/MHC-I | Hashtag_4 | M1/42; 30-F11; | BioLegend | 0.1-0.5µg/million cells |
| CD45/MHC-I | Hashtag_5 | M1/42; 30-F11; | BioLegend | 0.1-0.5µg/million cells |

Finally, samples were washed three times with PBS 2% of BSA and resuspended in PBS 2% BSA containing propidium iodide at dilution of 1:500. T and B cells were sorted using a BD FACS Aria III. After cell sorting, samples from different mice were pooled together and loaded per channel onto the Chromium controller, for generation of gel bead-in-emulsions. cDNA sequencing libraries were prepared using Single-cell 5’ reagent kits (10X Genomics) and converted using the MGIEasy Universal Library Conversion Kit (BGI) for sequencing on a MGISEQ-2000 instrument (BGI).

### CRISPR/Cas9 gene editing in naïve PbTII cells

Spleens from the C57BL/6J PbTII.Nzegfp donor mouse were collected and homogenized through a 70-μm cell strainer. Cells were lysed with Lysing Buffer (BD) and CD4^+^ PbTII GFP^+^ cells were enriched using CD4 (L3T4) microbeads (Miltenyi Biotec). To perform CRISPR/Cas9 knockout in naïve PbTII GFP^+^ cells, single guide RNAs (sgRNA) targeting Cxcr3 (5′-GAACAUCGGCUACAGCCAGG-3′,5′-UGAGGGCUACACGUACCCGG-3′), Cxcr5 (5′-UACCCACUAACCCUGGACAU-3′,5′-AGAGAAGGUCGGCUACUGCG-3′), and Cd19 (5′-AAUGUCUCAGACCAUAUGGG-3′) were purchased from Synthego (CRISPRevolution sgRNA EZ Kit, Synthego). To form sgRNA/Cas9 RNPs, 0.3 nmol sgRNAs were mixed with 0.6 μl Alt-R S.p. Cas9 nuclease V3 (10mg/ml, Integrated DNA Technologies) and incubated for 10 min at room temperature. Enriched naïve PbTII GFP^+^ cells were resuspended in 20μl of P3 Primary buffer (P3 Primary Cell 4D-Nucleofector X Kit, Lonza) and mixed with sgRNA/Cas9 RNP. Subsequently, cells were electroporated on the Lonza 4D-Nucleofector System (DN100), rested in the RPMI cell culture for 10min at 37°C, 5% CO_2,_ and then washed with PBS. Immediately, 3.5-1×10^5^ edited-PbTII GFP^+^ cells were adoptively transferred to each C57BL/6J mouse via *i.v.* injection. 5 days after adoptive transfer, mice were infected with *Pc*AS via lateral tail vein injection.

### Collection of tissues and *Slide-seqV2*

Spleen samples were placed in the cryomold immediately after dissection, covered in OCT medium, frozen on dry ice and stored at -80°C. Frozen samples were shipped to the Broad Institute of Harvard and MIT (Cambridge, MA, USA) and the *Slide-seqV2* was performed as described elsewhere (Stickels 2021). Briefly, fresh frozen tissue was cut into 5μm-thick sections and placed on a 3mm diameter circular array of 10μm diameter mRNA capture beads, referred to here as spots. Spots are coated in mRNA capture probes consisting of a location-specific barcode, a probe-specific UMI, and an mRNA-capturing polyT sequence. The tissue is permeabilised, mRNAs captured and ligated to probes, reverse transcribed, and then sequenced [31, 32]. Reads were aligned with STAR against GRCm38.81 and *Plasmodium chabaudi chabaudi* AS genome build 43 [67].

### Analysis of *Slide-seqV2* datasets

*Slide-seqV2* datasets were initially imported into R v3.5.1 and stored as Seurat v3 objects [35, 68]. To remove spots that were low quality, spots that fell outside of the circular *Slide-seqV2* arrays were removed via dbscan v1.1 clustering with an epsilon radius of 50 pixels (1 pixel = 0.665μm) [69]. To prepare the datasets for deconvolution, we removed irrelevant *Plasmodium chabaudi* AS mRNAs and filtered spots containing less than 100 UMIs. We selected array 4 from day 0 and array 1 from day 7 as representative samples for figures based on favourable combinations of high mRNA capture and minimal distortion to tissue microarchitecture during handling.

Subsequent analysis took place in R v4.2.0 and Seurat v4. Library size normalisation via Seurat’s NormalizeData() function was used to transform gene expression data for plotting. Because library size varies across biologically distinct tissue regions, we performed a neighbourhood-based library size normalisation in which each spot’s gene expression is adjusted so the total number of UMIs is equal to the average UMI count for all spots in a 75μm radius. The adjusted gene expression values were then transformed via addition of 1 pseudocount followed by natural log transformation using R’s log1p function. This was inspired by stlearn [36].

### Local neighbourhood averaging

Unbiased clustering based on spot-specific gene expression does not consider the gene expression of nearby spots and thus does not use spatial information. To incorporate spatial information, we used an approach we termed local neighbourhood averaging (LNA), inspired by stlearn [36]. In this approach, we performed principal component analysis based on neighbourhood-adjusted library size-normalised gene expression generated as above. We then averaged principal component scores so that each spot’s score for each principal component is the mean of the principal component score of all spots within a 50μm radius. These averaged principal component scores thus incorporated information from nearby spots and could be used as input for unbiased clustering or UMAP.

### Analysis of scRNA-seq datasets

Whole splenocyte scRNA-seq data were aligned via Cell Ranger v3.1.0 against mouse genome v2020-A supplied by 10X Genomics. Data were initially analysed via R v3.5.1 and Seurat v3. Doublet removal was performed with DoubletDecon v1.1.4 and automated annotation was performed with SingleR [70, 71]. Downstream analysis took place in R v4.2.0 and Seurat v4. We additionally used scanpy v1.9.1 in python v3.9.6 and reticulate v1.25 in R v4.2.0 for transferring dataset between Seurat and AnnData formats [72].

Transcriptome and V(D)J sequences of PbTII cells and polyclonal activated CD4^+^ T cell enriched by CXCR3 and CD11a expression for antigen experience were aligned via Cell Ranger 3.1.0 (10X Genomics) against mouse genome v2020-A supplied by 10X Genomics. Transcriptome of splenic CD19^+^ B220^hi-int^ B cells were also aligned via Cell Ranger 3.1.0 (10X Genomics) against mouse genome v2020-A supplied by 10X Genomics. Pre-processing of the data was performed using Seurat v4 and R v4.2.0. Doublets were excluded based on hashtag expression. For the B cell dataset, low quality cells were removed low-quality cells outside the threshold of 200-4000 expressed genes and those with mitochondrial gene content >15% were removed. T cells were removed based on high mitochondrial content (>10%) and low or anomalously high total gene expression (<300 or >6000). For the B cell dataset, differential gene expression analysis was performed using the FindMarkers() function in Seurat v4 with Wilcoxon rank sum test used to identify statistically significant differentially expressed genes. Geneset enrichment analysis (GSEA) was performed against biological processes GO terms using WebGestalt [73].

### Cell type annotation

After performing quality control of the whole spleen scRNA-seq dataset (Supp. Methods Fig. 1A), we embedded the single cell transcriptomes via PCA and UMAP (Supp. Methods Fig. 1B). We next attempted to define which cell types were captured at each timepoint by examining differences in gene expression across the dataset (Supp. Methods Fig. 1C), and with an automated annotation tool, *SingleR*, which assigns cell type labels based on a reference dataset (Supp. Methods Fig. 1D). Some cell types such as endothelial cells (6), fibroblasts (1), and dendritic cells (2) were inferred at low frequency by *SingleR*, which we ignored as around 25 cells are required to form a cell type reference for deconvolution. We noted the majority of cells appeared to be lymphocytes, expressing either *Cd3e, Ms4a1*, which we defined as T and B cells respectively (Supp. Methods Fig. 1D-E). We also noted *Sdc1* expression, indicating antibody-secreting cells, which we grouped with the B cells. We next examined gene expression among other subsets that *SingleR* had identified such as NK cells, erythrocytes, granulocytes, monocytes, and macrophages. We noted a cluster lacking *Cd3e* but positive for *Klrg1*, which we inferred were NK cells. We also noted a cluster lacking most other markers but positive for *Hba-a1*, a haemoglobin gene, which we inferred were erythrocytes. Lastly, we noted a complex group of myeloid cells variously expressing genes such as *Itgam, Itgax, Cd14*, and *Ngp* (Supp. Methods Fig. 1C)*. SingleR* inferred a group of granulocytes which we noted matched with *Ngp*, a neutrophilic granule protein gene, thus we annotated these as granulocytes. Finally, we labelled the other myeloid cells as a single cluster for further analysis (Supp. Methods Fig. 1E).

B cells, myeloid cells, and T cells all occur in a variety of functional and anatomical subsets with relevance to malaria. Thus, to identify which subsets we had captured, we embedded, B cells (Supp. Methods Fig. 2A), myeloid cells (Supp. Methods Fig. 2D), and T cells (Supp. Methods Fig. 3A) via PCA and UMAP. We first began annotating B cells and noted that cells collected at day 0 or day 7 were segregated via UMAP embedding, which we speculated was due to activation and differentiation. Genes such as *Fcer2a*, encoding CD23, and *Cr2*, encoding CD21, revealed follicular and marginal zone B cells (Supp. Methods Fig. 2B-C), whereas *Zbtb20* revealed B1 cells (Supp. Methods Fig. 2B-C). These account for most B cells captured at day 0, save for some antibody-secreting cells which were identified by *Xbp1* (Supp. Methods Fig. 2B-C). At day 7, we noted two activated subsets based on *Mki67*, which we inferred were antibody-secreting plasmablasts based on *Xbp1* or pre-germinal centre cells based on lack of this gene. Some cells were neither plasmablast nor pre-germinal centre cells, which we inferred were follicular cells. However, they differentially upregulated *Socs3* among other genes at high levels compared to day 0 follicular cells. Because there were no other follicular cells at day 7, and these cells were polyclonal, we reasoned that these must be cells with antigen receptors irrelevant to malaria undergoing a bystander response. Thus, we defined plasmablasts (ASCs), pre-germinal centre, and bystander cells at day 7 (Supp. Methods Fig. 2B-C).

We next annotated myeloid cells. After embedding via UMAP (Supp. Methods Fig. 2D), we examined gene expression to determine which subsets were captured. We first examined gene expression, including monocyte markers such as *Itgam, Ly6c2*, cDC markers such as *Itgax*, *H2-Ab1* (MHC-II subunit)*, Cd4*, and *Cd8b1*, and a pDC marker *Siglech* (Supp. Methods Fig. 2E). Furthermore, we noted a population of *Ms4a1* (encoding CD20)-expressing cells which were removed as doublets. pDC, cDC1, and cDC2 cells were evident based on *Siglech, Cd8b1*, and *Cd4*, respectively (Supp. Methods Fig. 2F); all other cells at day 0 were labelled as monocytes. In addition, at day 7, a subset of monocyte-like cells lacking *Itgam* and *Ly6c2* were labelled as macrophages (Supp. Methods Fig. 2F), while remaining cells expressing these genes were annotated as monocytes. All of the cells we annotated as monocytes or macrophages expressed associated phagocytic genes such as *Cd68, Lamp1*, and *Lamp2*, and the Fc receptor *Fcgr1*, although low samples numbers meant the subsets were difficult to resolve.

Finally, we performed detailed annotation of T cells. After embedding T cells via PCA and UMAP (Supp. Methods Fig. 3A), we attempted to distinguish CD4^+^ and CD8^+^ T cells by expression of *Cd4* and *Cd8b1*. Because there are two CD8 genes, *Cd8a* and *Cd8b1*, we reasoned that cells lacking both genes were likely CD4^+^ cells, which were labelled as such while the rest were annotated as CD8^+^ (Supp. Methods Fig. 3B). One aberrant cluster was defined by expression of both T cell markers and B cell markers such as *Ms4a1*, encoding CD20, and was removed (Supp. Methods Fig. 3A-B). CD4^+^ T cells were re-embedded and annotated according to expression of marker genes (Supp. Methods Fig. 3C-E). Naïve and bystander cells were identified by expression of *Ccr7* and lack of differentiation markers such as *Cxcr6* or *Cxcr5.* Tregs were identified by *Foxp3* expression, innate-like natural killer T cells via *Zbtb16* [74], and a pool of what we speculated were pre-existing memory T cells by slightly reduced *Ccr7* and increased *Cxcr5*, although we excluded these cells for deconvolution due to lack of other distinguishing markers, and because memory-like cells were captured in the day 7 polyclonal CD4^+^ T cell dataset. Finally, we identified what were likely malaria-responding cells by *Mki67* for proliferating cells, *Cxcr5* for Tfh, and *Cxcr6* for Th1 (Supp. Methods Fig. 3C-E). Lastly, to identify CD8^+^ T cell subsets, we re-embedded the data from (Supp. Methods Fig. 3B) and identified naïve, bystander, and activated subsets by expression of *Ccr7* compared to markers of cytotoxicity such as *Klrg1* and *Gzmb* (Supp. Methods Fig. 3F-H).

### Cell type deconvolution

*Slide-seqV2* datasets were deconvoluted with cell2location v0.1 in python v3.10.4 and spacexr v2.0.0, containing the RCTD algorithm, in R v4.2.0 [37, 38].

Both algorithms require a single cell transcriptomic reference to deconvolute spatial arrays. As a reference, we used the immune cell datasets described above as well as a published splenic stromal cell dataset from which we selected fibroblastic reticular cells, follicular dendritic cells, red pulp reticular cells, and adventitial reticular cells as most relevant and abundant [48]. To deconvolute day 7 pucks, we used a combination of the whole spleen scRNA-seq dataset with the polyclonal CD4^+^ T cell dataset as the latter had greater numbers of Th1, Tfh, Tcm, and iTreg cells. We excluded Th1, Tfh, and Tcm cells from the whole spleen scRNA-seq dataset but included subsets that the CXCR3^+^CD11a^hi^ gating strategy did not capture, including bystander cells.

RCTD was run simultaneously on all datasets collected at the same timepoints via the run.RCTD.replicates() function in doublet mode. For spot class estimation, that is, determining which spots are a single cell type or mixtures of two known or unknown cell types, we combined similar cell types into groups. We specified stromal; B cell; T and NK cell; myeloid; and erythrocyte classes in both timepoints.

cell2location was also run on all day 0 datasets simultaneously and all day 7 datasets simultaneously, treating each array as a batch. Genes expressed in less than 500 spots across all datasets were discarded from spatial datasets. The reference model was trained for 1000 epochs with lr=0.01. The cell2location model was trained for 500 epochs with lr=0.01, N_cells_per_location=1, and detection_alpha=200.

### Cell-cell colocalisation inference

To determine whether cell types occupied similar spatial locations, we defined spatial neighbourhoods with a radius of 50μm around all beads. Inside these spatial neighbourhoods, we averaged the abundance of each cell type, inferred by *RCTD or cell2location*, across all beads in the neighbourhood. We then computed Spearman correlations to determine cell type pairs whose abundances correlated in these neighbourhoods.

### Cell-cell interaction inference

We inferred cell-cell interactions using CellChat v1.6.1 in R v4.2.0 [44]. We followed the default workflow and settings for inferring cell-cell communication in a single dataset, although in *computeCommunProb* type=’truncatedMean’ was used to avoid false negatives. Interactions with p > 0.05 were filtered.

### Plotting

We used R 4.2.0 packages ggplot2 v3.3.6 [75] and cowplot v1.1.1, as well as prism v9, for plotting. Figures were generated in Adobe Illustrator and schematics were generated with Biorender.

When plotting deconvolution output from *cell2location*, we normalised so that the total cell type content at each bead was equal to 1, as opposed to an estimated number of nuclei. When plotting stromal and CD4^+^ T cell subsets, we condensed the colour scales so that any bead with a proportion of these cell types greater than 50% of maximum in the dataset appeared as the highest value in the colour scale. This transformation was not applied when performing cell-cell colocalisation inference (above).

## Acknowledgements

This research was supported by The University of Melbourne’s Research Computing Services, the Petascale Campus Initiative, and the LIEF HPC-GPGPU Facility hosted at the University of Melbourne [76]. This Facility was established with the assistance of LIEF Grant LE170100200. We thank the Biological Research Facility at the Peter Doherty Institute as well as the Doherty Institute node of the Melbourne Cytometry Platform for facility access and technical assistance. We acknowledge Prof DI Godfrey, University of Melbourne, for provision of CD1d tetramer for NKT cell exclusion during flow cytometric assessments. This work was partly funded by grants awarded to AH by the Australian National Health & Medical Research Council: Ideas Grant Numbers 2010784 & 1180951.

## Extended Data

**Extended Data Figure 1.**
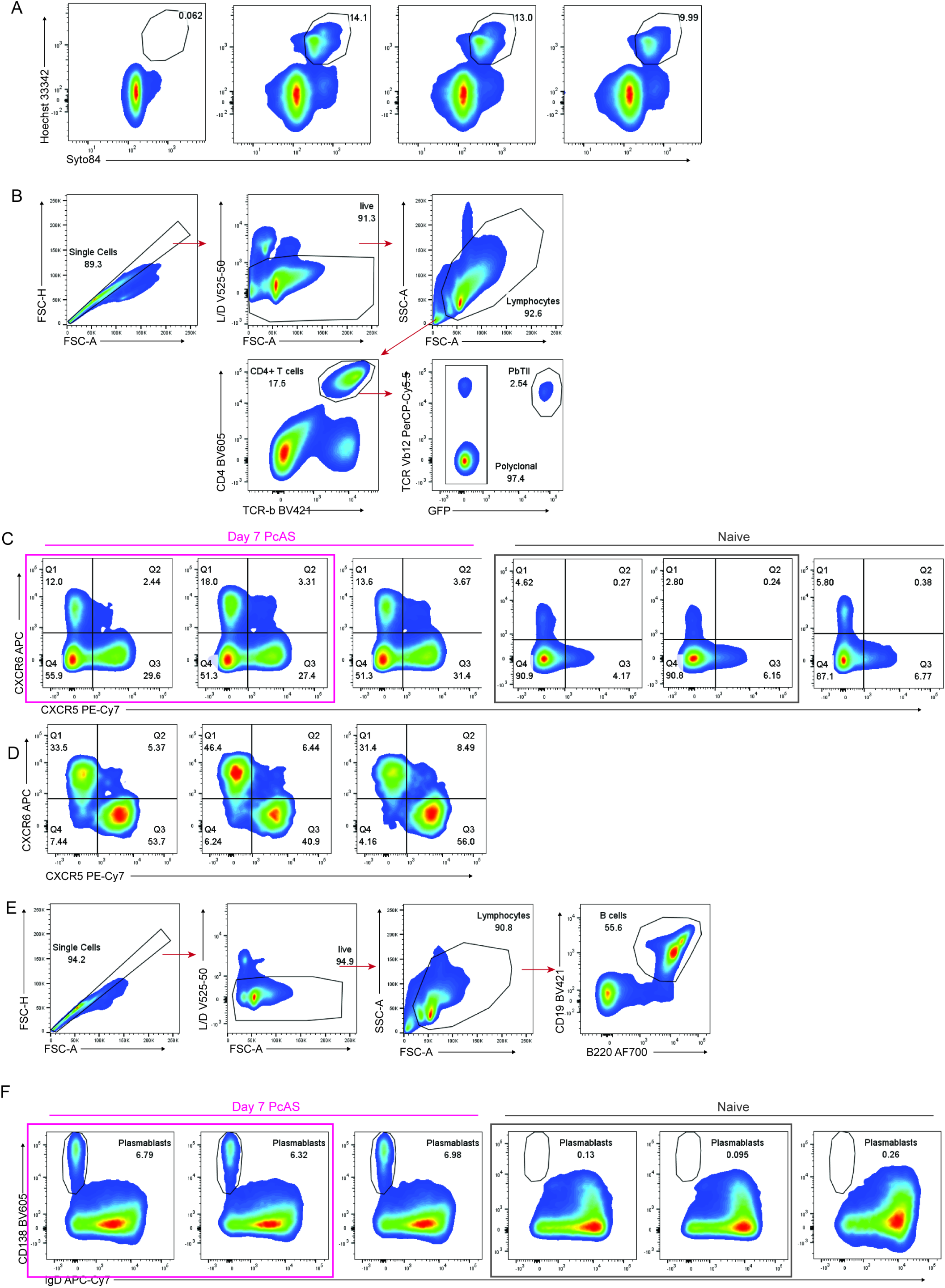
Quality control of CD4+ T cell and B cell responses to *Pc*AS infection in mouse samples subjected to *Slide-seqV2.* (A) Parasitemia measurements in one naïve and three infected mice on day 7 p.i. (B) Representative FACS plots showing gating strategy to analyse polyclonal and PbTII CD4 T cells in naïve and infected mice at 7 days p.i. (C) FACS plots showing CXCR5 *versus* CXCR6 expression by polyclonal CD4^+^ T cells in naïve and infected mice at 7 days p.i. The infected and I samples used for SlideSeq-V2 are highlighted in pink and grey, respectively. (D) FACS plots showing CXCR5 *versus* CXCR6 expression by PbTII cells in infected mice at 7 days p.i. (E) Representative FACS plots showing gating strategy to analyse B cells in naïve and infected mice at 7 days p.i. (F) FACS plots showing IgD *versus* CD138 expression by B cells, for plasmablasts gating, in naïve and infected mice at 7 days p.i. The infected and naive samples used for SlideSeq-V2 are highlighted in pink and grey, respectively.

**Extended Data Figure 2.**
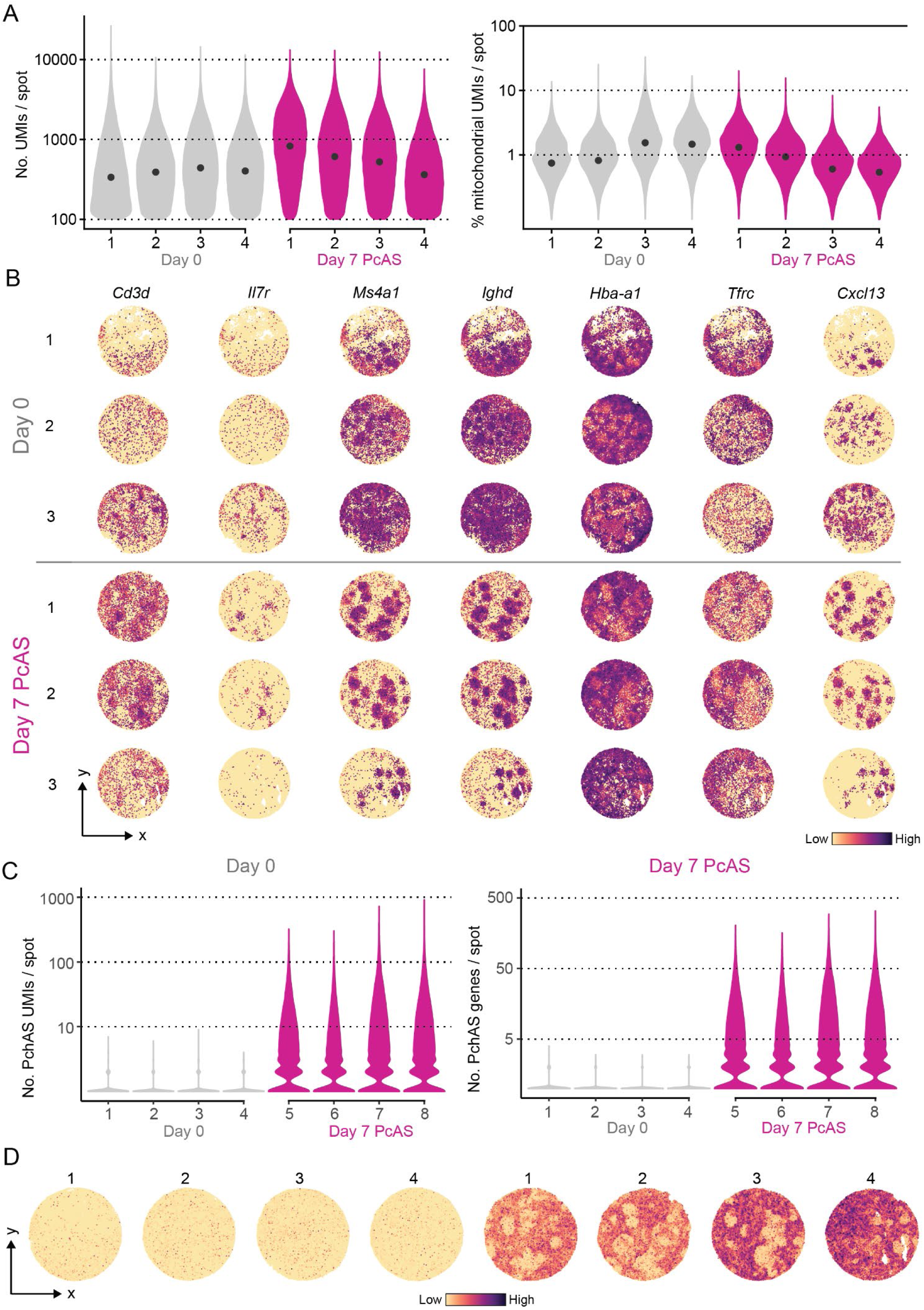
Quality control and initial assessment of *Slide-seqV2* arrays. (A) Violin plots of UMIs per spot (left) and percent mitochondrial content per spot (right) in day 0 and day 7 *Slide-seqV2* arrays. (B) Violin plots of number of *Plasmodium chabaudi chabaudi* AS-derived UMIs (left) and genes (right) detected per spot per array. (C) Normalised expression of spatially confined genes indicating T cells (*Cd3d*, *Il7r*), B cells (*Ms4a1, Ighd*), red blood cells (*Hba-a1, Tfrc*) and stromal cells (*Cxcl13*) in both uninfected and *Pc*AS spleens in pucks not shown in Fig. 1C. (D) Detection of *Plasmodium chabaudi chabaudi AS* mRNAs per spot in each array, mapped to spatial coordinates.

**Extended Data Figure 3.**
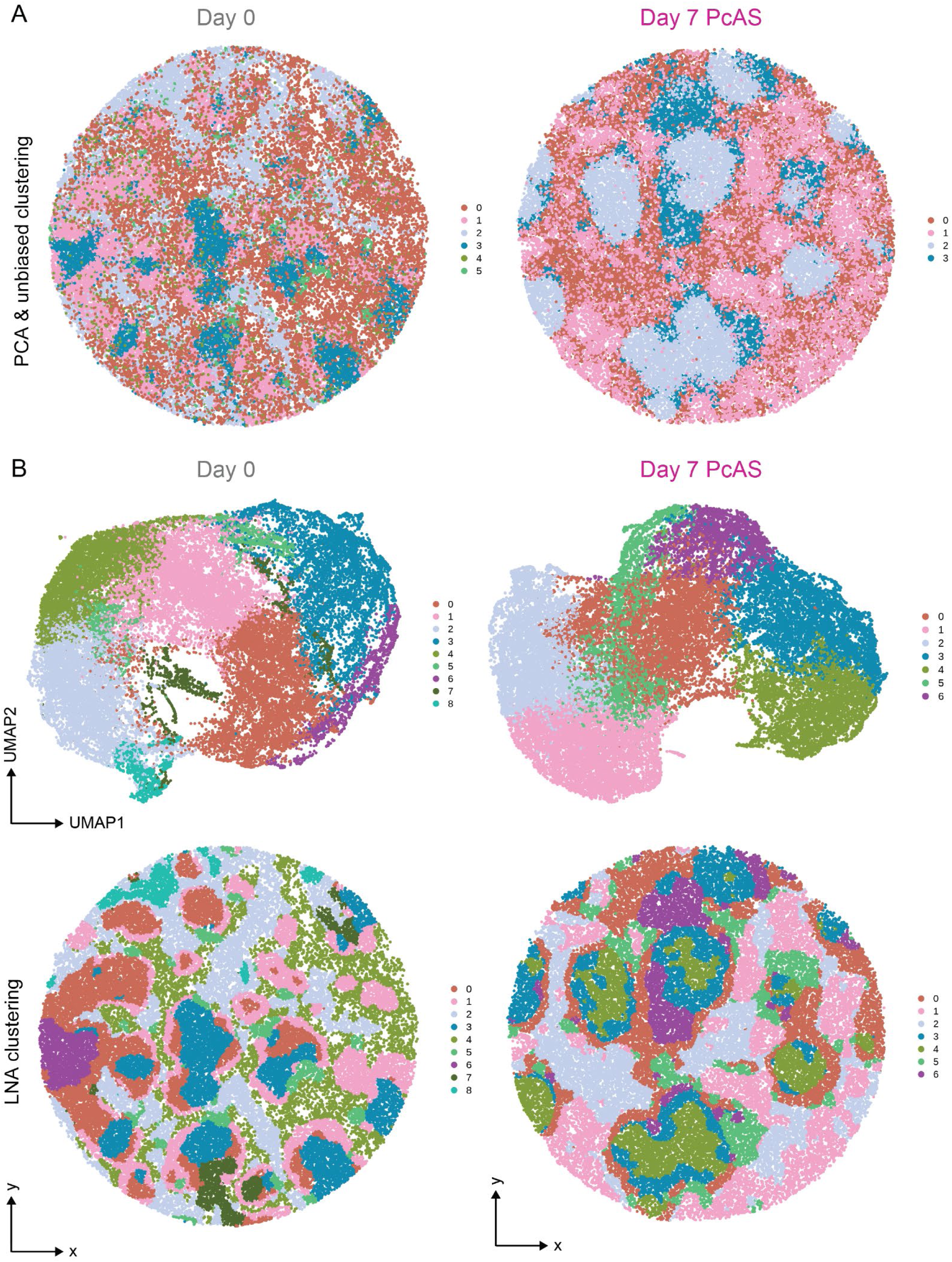
Local Neighbourhood Analysis reveals spatially confined microanatomical niches. (A) Unbiased clustering of *Slide-seqV2* beads in representative day 0 (left) arrays based on standard Principal Component Analysis (top) in representative day 0 (left) and day 7 (right) pucks, compared to unbiased clustering based on Local Neighbourhood Analysis-based clusters (bottom) in same pucks. (B) LNA-based UMAP embeddings of beads from day 0 (left) and day 7 (right) pucks shown below.

**Extended Data Figure 4.**
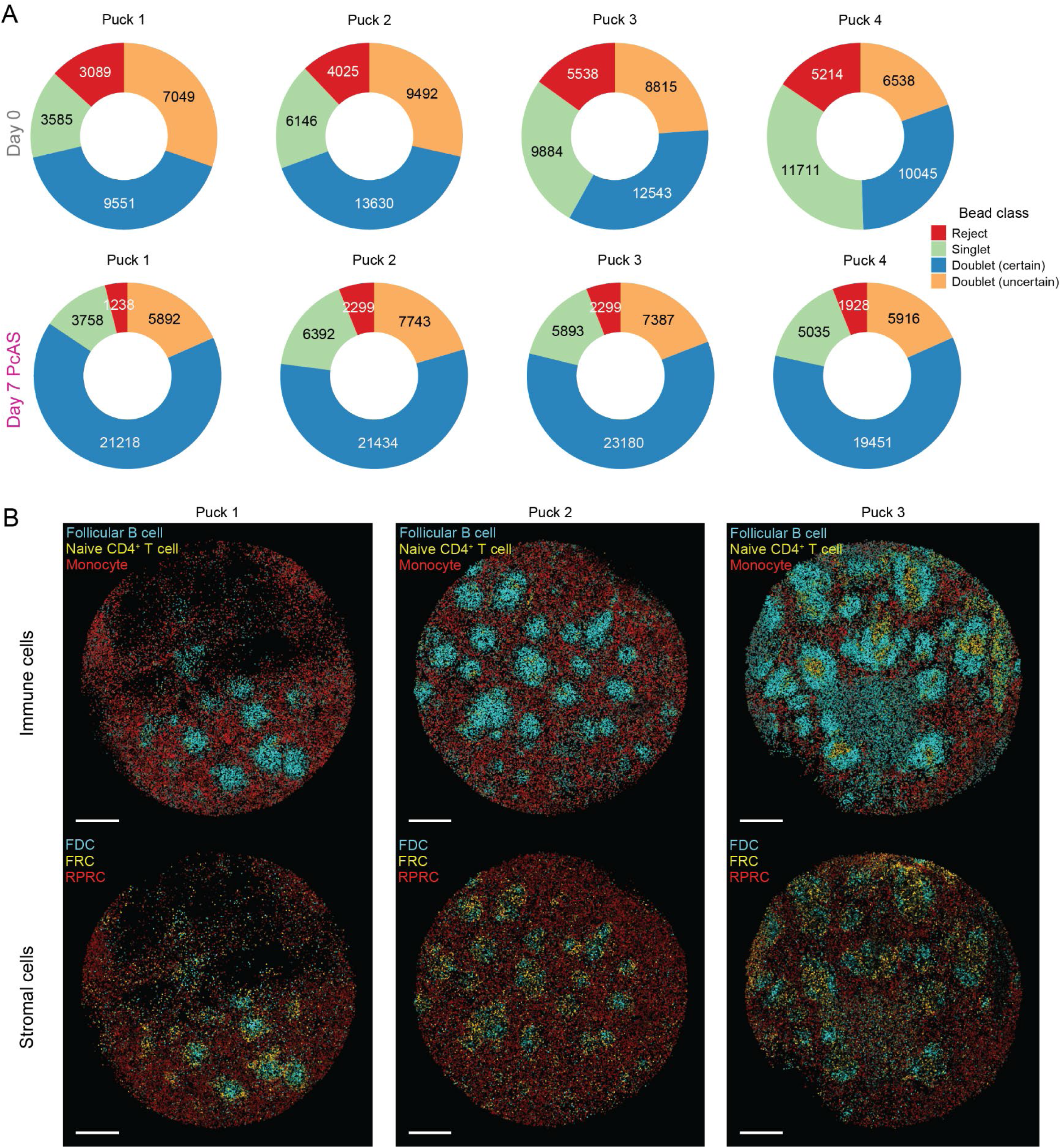
Integration of scRNA-seq and *Slide-seqV2* data via *RCTD* reveals cell type locations in naïve mouse spleen. (A) Proportion of beads in each day 0 (top) and day 7 (bottom) puck inferred by *RCTD* to consist of a single cell type (singlet); two known cell types (doublet (certain)); a known cell type and unknown cell type (doublet (uncertain)); and unknown cell types (reject). Absolute numbers displayed on each sector. (B) Top: Follicular B cells (cyan), CD4^+^ T cells (gold), and monocytes (red) inferred by *RCTD* in other day 0 *Slide-seqV2* pucks. Bottom: locations of corresponding stromal cell subsets. FDCs (cyan), FRCs (gold), and RPRCs (red). Stromal cell signatures plotted on condensed scale for visibility. Scale bar shown is 500µm.

**Extended Data Figure 5.**
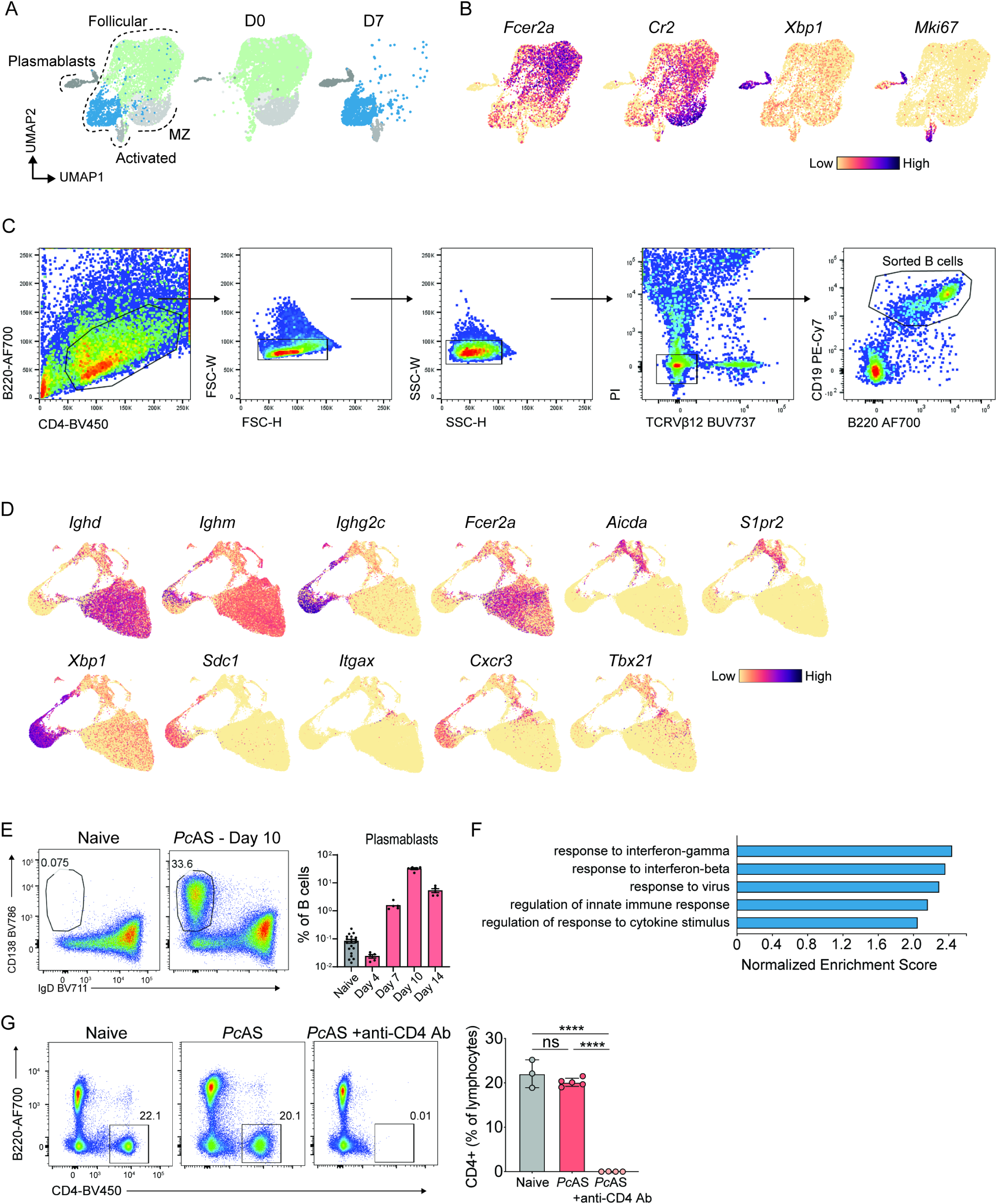
Quality control of scRNA-seq analysis of B cells and bystander B cell assessment from *Pc*AS infected mice. (A) High dimensional clustering showing follicular (coloured) and mature-like B cells (ASCs, pre-GC, B1 and MZ B cells) (grey) represented by the combined UMAP and UMAPs split by timepoints (D0 and D7). (B) Feature plots of genes associated with follicular B cells (*Fcer2a*), marginal zone B cells (*Cr2*), ASCs (*Xbp1*), pre-GC (*Mki67*) and B1 cells (*Zbtb20*). (C) FACS gating strategy used to sort total CD19^+^ B220^hi-int^ B cells used in the scRNA-seq experiment. (D) Violin plots showing scRNA-seq QC metrics number of molecules, number of UMIs and percentage of mitochondrial RNA detected per cell, split across time points post *Pc*AS infection. (E) Ridge Plots of the hashtag deconvolution used to identify individual mouse samples in the scRNA-seq dataset. (F) Feature plots of genes associated with naive, follicular B cells (*Ighd, Fcer2a*), isotype-switched B cells (*Ighm, Ighg2c*), germinal centre B cells (*Aicda, S1pr2*), plasmablasts (*Xbp1, Sdc1*) and atypical B cells (*Itgax, Cxcr3, Tbx21*). (G) Representative FACS plot of day 10 plasmablasts (CD138^+^, IgD^−^ B cells) and bar graph of frequency of plasmablasts relative to all B cells in naïve and days 4, 7, 10 and 14 post *Pc*AS infection. n=20 biological replicates for naïve, n=5 biological replicates for each timepoint post *Pc*AS infection, data representative of three independent experiments. (H) Gene set enrichment analysis of the differentially expressed genes from D4 bystander cluster versus D0 naive cluster, showing the top 5 significantly enriched biological processes GO terms. (I) Representative FACS plots and bar graph of CD4 expression on lymphocytes 7 days post PcAS infection after CD4 T cell depletion. Statistical test used is one-way ANOVA with Tukey’s multiple comparisons test, n=3-5 biological replicates. Data representative of two independent experiments. Data are ± S.D. p-value **** <0.0001.

**Extended Data Figure 6.**
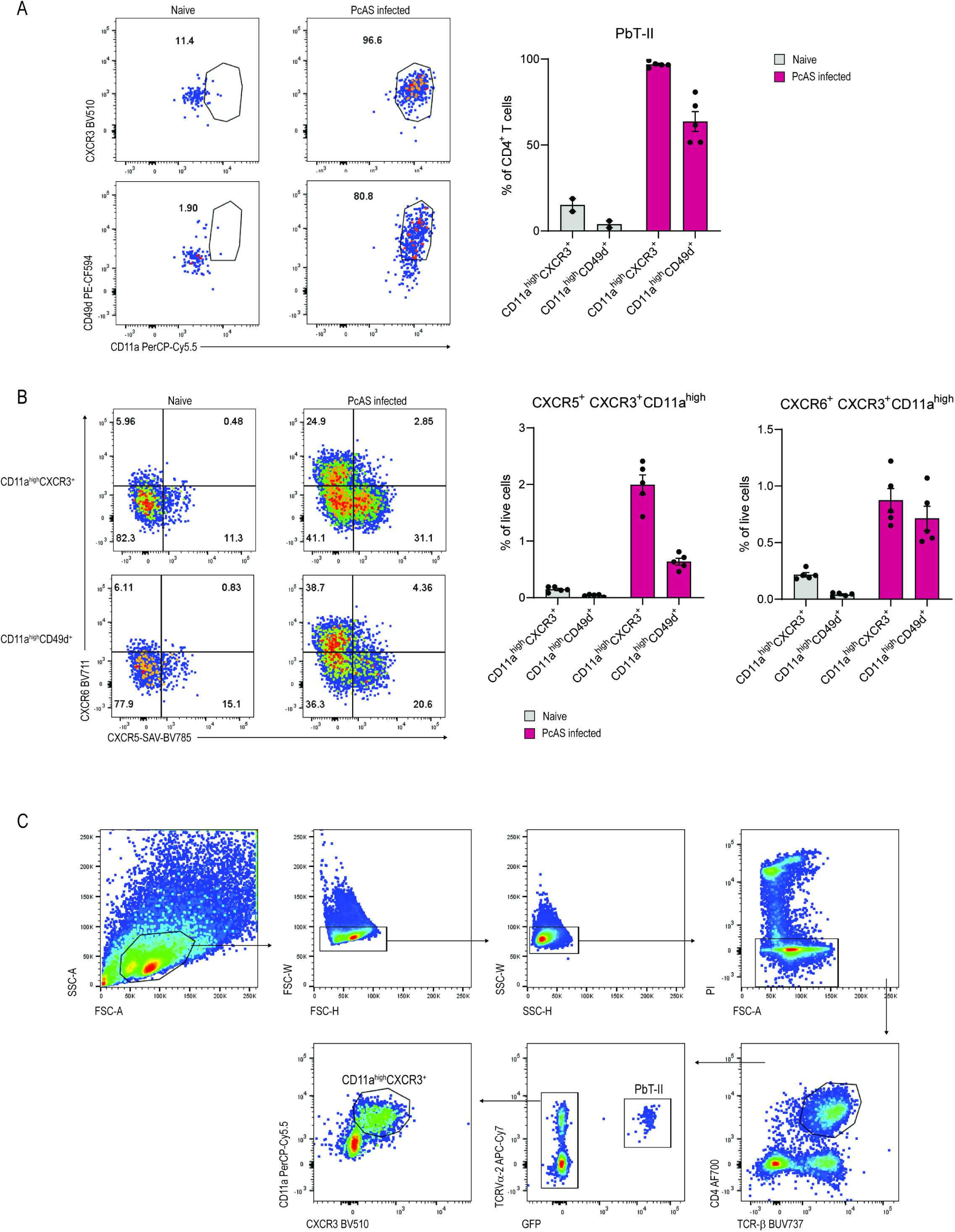
CXCR3 expression in association with CD11a captures a large proportion of CD4+ T cells activated during *Pc*AS infection. (A) Representative FACS plots and bar graph showing the expression of CD11a, CXCR3 and CD49d by PbTII cells in naïve and infected mice 7 p.i., data representative of three experiments (5 mice per group, per experiment). (B) Representative FACS plots and bar graphs showing the expression of CXCR6 and CXCR5 by CD11a^high^CXCR3^+^ and CD11a^high^CD49d^+^ CD4^+^ T cells in naïve and infected mice on day 7 p.i. Data representative of three experiments (5 mice per group, per experiment). (C) FACS gating strategy used to isolate PbTII and CD11a^hi^CXCR3^+^ cells used in the scRNA-seq experiment.

**Extended Data Figure 7.**
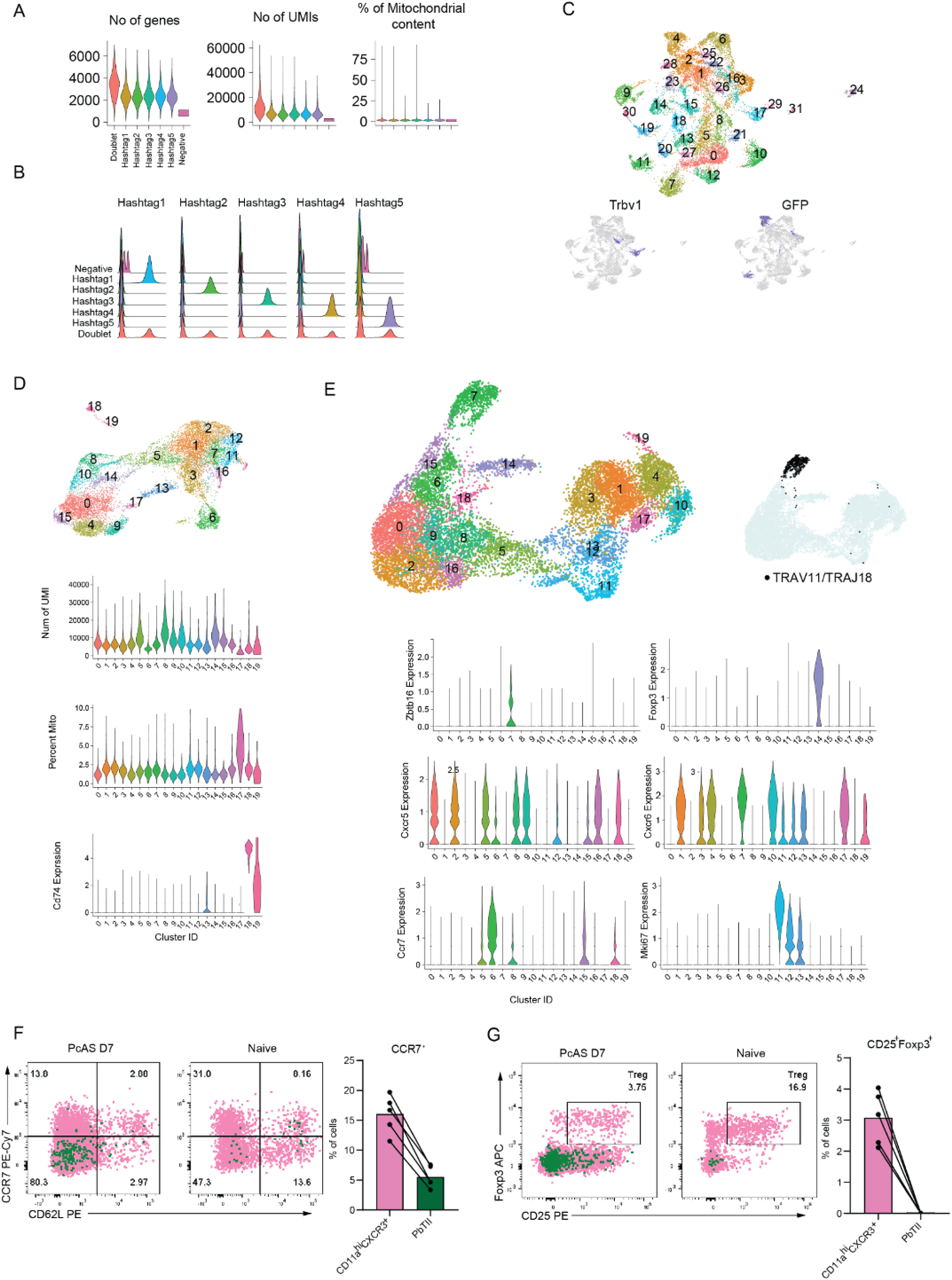
Quality control of scRNA-seq assessment of PbTII and CD11a^hi^CXCR3^+^ cells. (A) Violin plots showing scRNA-seq QC metrics number of UMIs, number of molecules and percentage of mitochondrial RNA detected per cell, split across hashtags. (B) Ridge Plots of the hashtag deconvolution used to identify individual mouse samples in the scRNA-seq dataset. (C) UMAP visualization of unsupervised clustering prior to the removal of GFP, *Trav/j* and *Trbv/j* genes from the gene expression matrix. (D) UMAP visualization of unsupervised clustering and violin plots of QC metrics used to identify and remove contamination with antigen presenting cells (clusters 18 and 19) from the dataset. (E) UMAP visualization of unsupervised clustering and Trav11/Traj18 expression, and Violin Plots of *Zbtb16, Foxp3, Cxcr5, Cxcr6, Ccr7* and *Mki67* used to identify and remove contamination with NKT cells. (F) Representative FACS plots of CCR7 and CD62L expression by PbTII (green) and CD11a^hi^CXCR3^+^ cells (pink) in naïve and and infected mice seven days p.i. with *PcAS*, and bar graphs showing frequencies of CCR7^+^ cells amongst CD11a^hi^CXCR3^+^ and PbTII cells in infected mice. Data representative of two experiments (5 mice per group, per experiment). (G) Representative FACS plots and bar graphs showing frequencies of Treg cells amongst CD11a^hi^CXCR3^+^ and PbTII cells in naïve and and infected mice seven days p.i. with *Pc*AS.

**Extended Data Figure 8.**
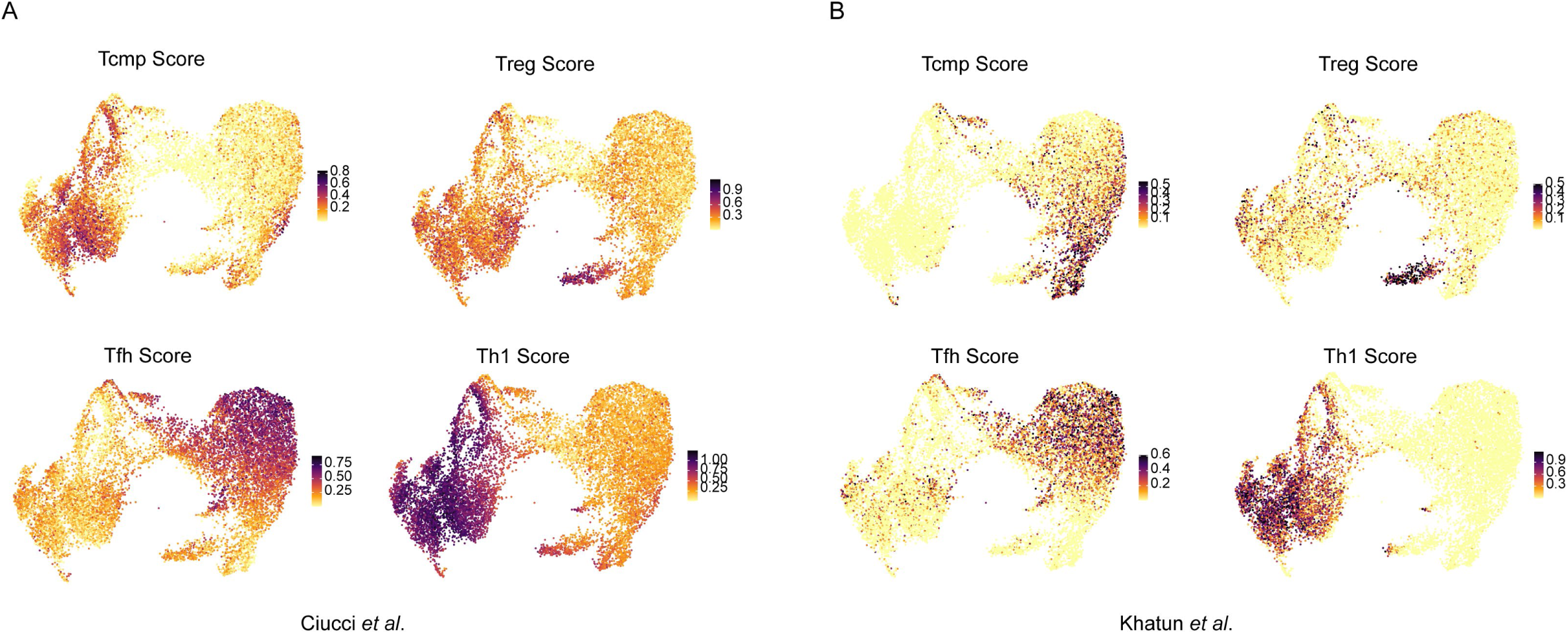
Cell-type module score UMAPS based on published lists of genes. UMAP representations CD11a^hi^CXCR3^+^ polyclonal and GFP^+^PbTII cells 7 days post *Pc*AS infection showing module scores for Tcmp, Treg, Tfh and Th1 cells calculated based on lists of genes published by (A) Ciucci *et al*. (2019) or (B) Khatun *et al*. (2021).

**Extended Data Figure 9.**
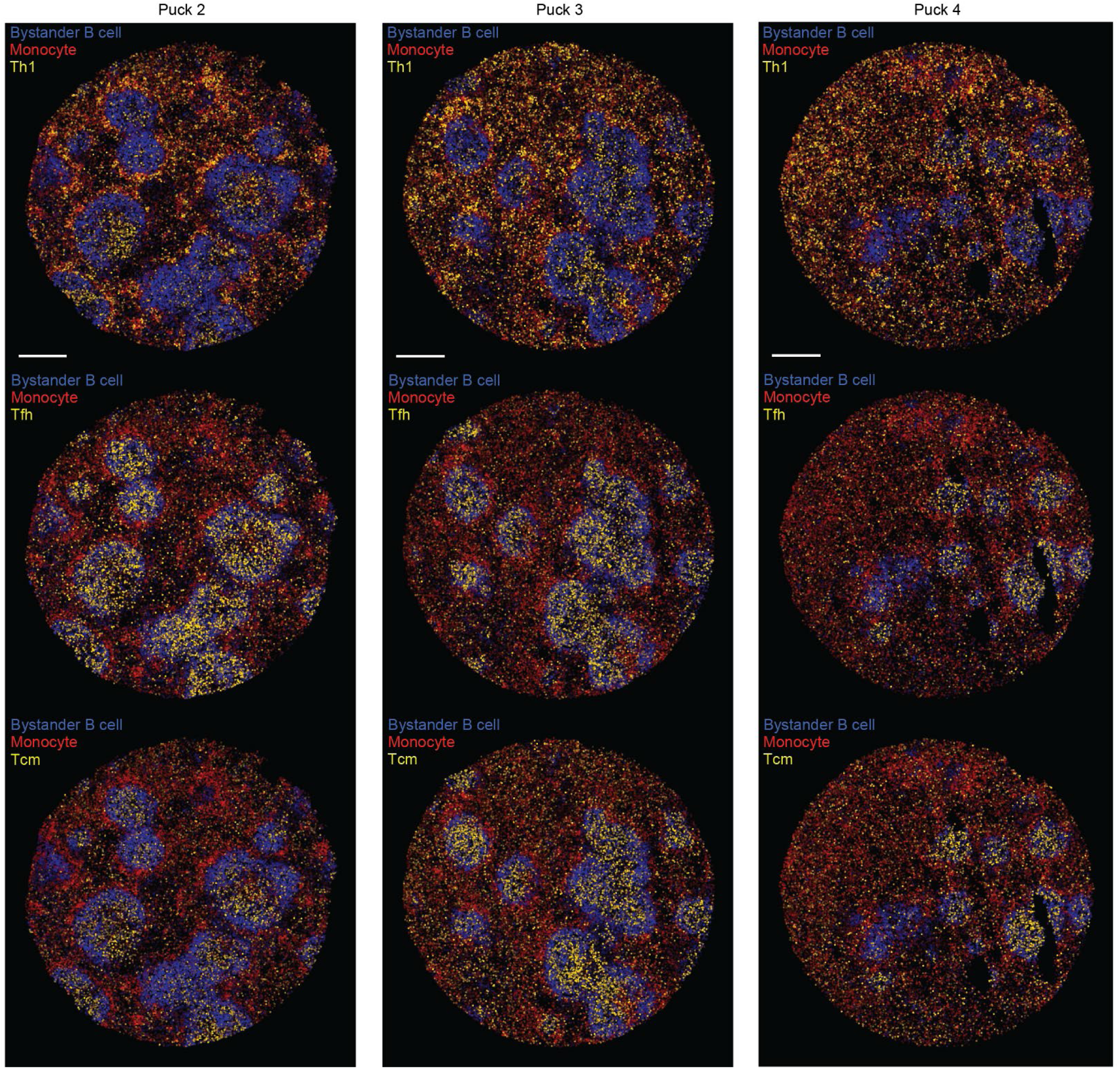
*RCTD* maps CD4^+^ T cell subsets during malaria infection. Bystander B cells (blue), monocytes (red), and CD4^+^ T cell subsets (top to bottom: Th1, Tfh, and Tcm, in gold) inferred by *RCTD* in day 7 *Slide-seqV2* pucks not shown in (Fig. 5B). CD4^+^ T cell signals doubled for visibility. Scale bar is 500µm. T cell signatures shown on shortened colour scale for visualization purposes.

**Extended Data Figure 10.**
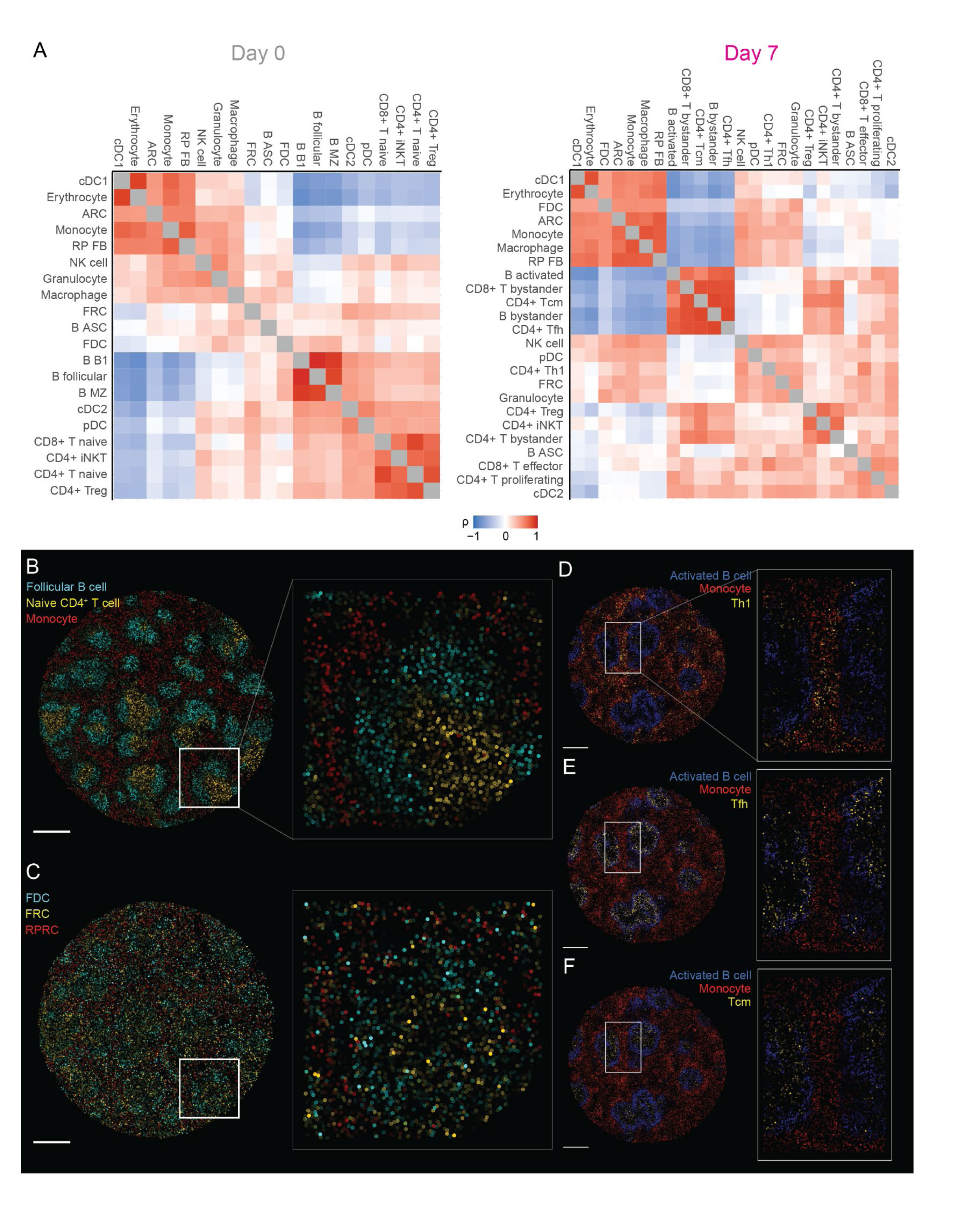
*cell2location* reveals cell type location locations before and during malaria infection. (A) Spearman correlations of all mapped cell type pairs at day 0 (left) and day 7 (right) in space. Based on locations inferred by *cell2location* and calculated over a neighbourhood of radius 50μm. (B) *Cell2location*-inferred locations of follicular B cells (cyan), naïve CD4^+^ T cells (gold), and monocytes (red) in a naïve day 0 spleen. Scale bar shown is 500μm. (C) *Cell2location*-inferred locations of FDCs (cyan), FRCs (gold), and RPRCs (red) in a naïve day 0 spleen. Scale bar shown is 500μm. Stromal cell signatures plotted on shortened colour scale for visualisation purposes. (D-F) *Cell2location*-inferred locations of activated B cells (blue), monocytes (red), and CD4^+^ T cell subsets (gold) including Th1 (D), Tfh (E), and Tcmp (F). Scale bar shown is 500μm. CD4^+^ T cell subsets plotted on shortened colour scale for visualisation purposes.

**Extended Data Figure 11.**
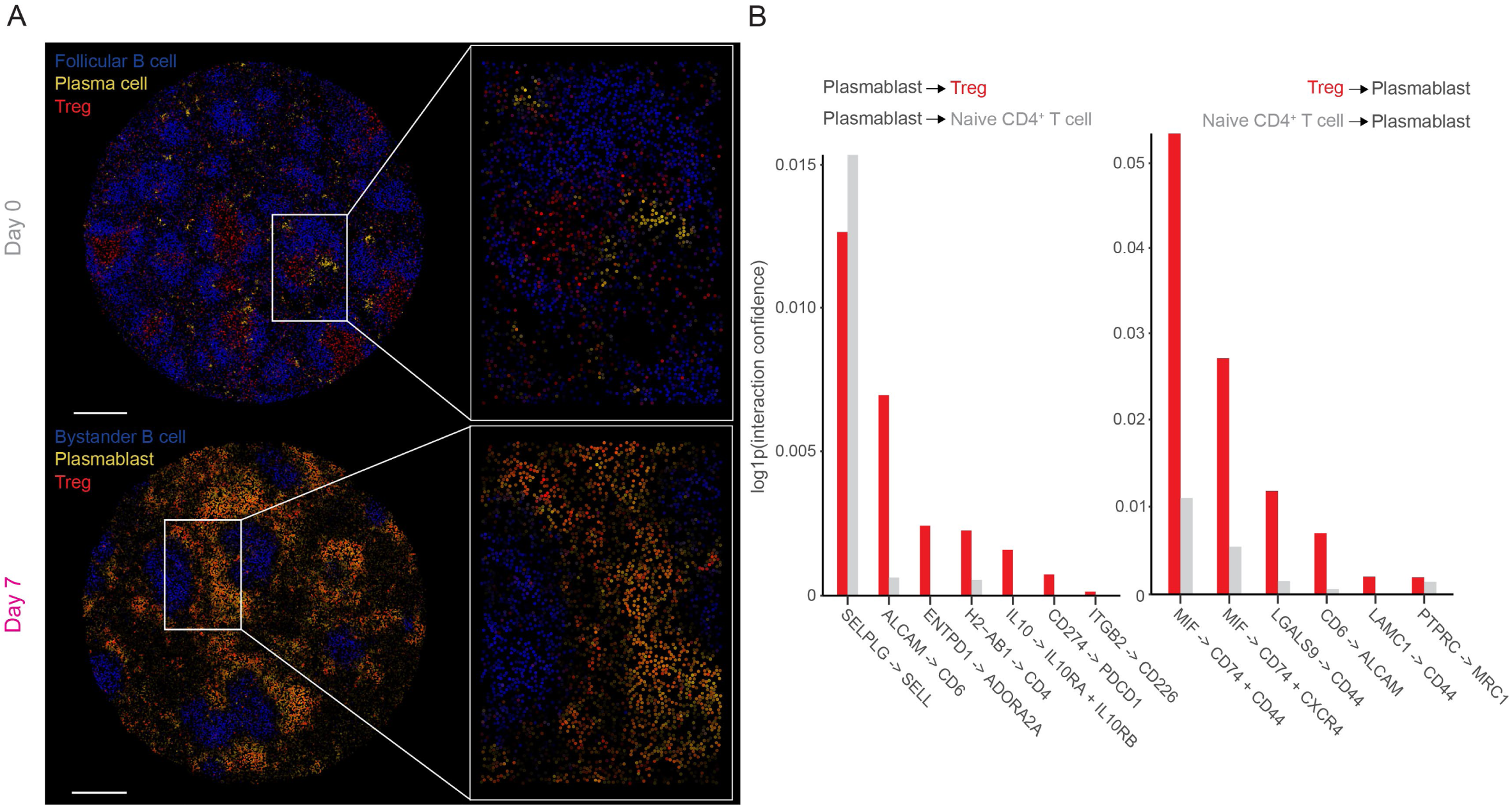
Plasmablasts and Tregs colocalise during malaria infection and express compatible ligand-receptor pairs. (A) Locations of antibody-secreting cells (i.e. plasma cells and plasmablasts) (red), regulatory T cells (cyan), and non-activated follicular B cells (dark blue) before (top) and during (bottom) malaria infection, inferred by *RCTD.* Scale bar shown is 500µm. (B) Ligand-receptor interactions from plasmablasts to Tregs (cyan) or to naïve CD4^+^ T cell (grey) controls (left), and from Tregs (cyan) or from naïve CD4^+^ T cell (grey) controls to plasmablasts (right) ranked by confidence

**Extended Data Figure 12.**
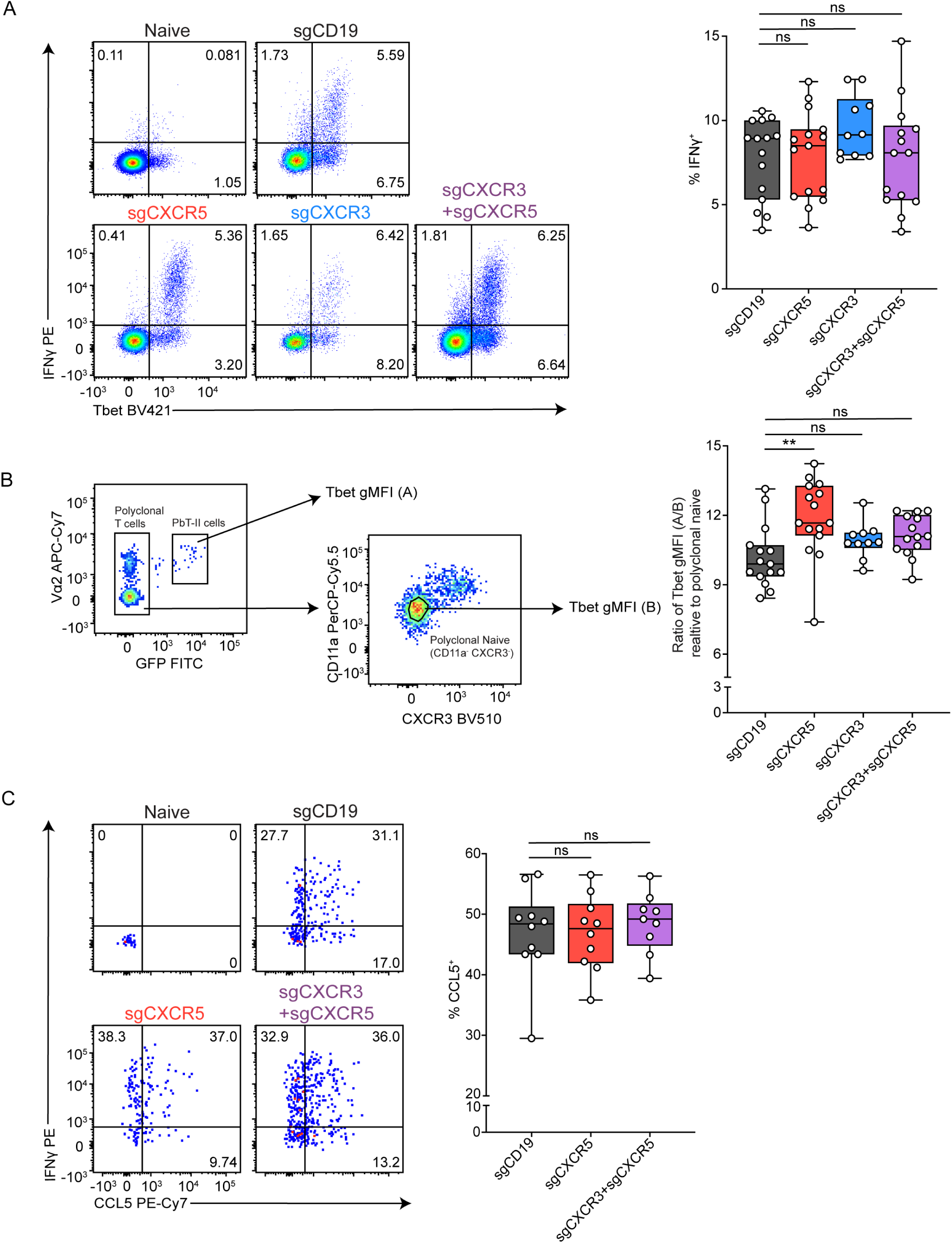
Effect of CXCR5-deficiency on Th1 differentiation. (A) Representative FACS plots and bar graph of direct ex vivo staining of IFN-γ^+^ in sgCD19, sgCXCR3, sgCXCR5, or sgCXCR3+sgCXCR5 polyclonal CD4^+^ T cells at day 7 p.i. (B) Gating strategy for Tbet geometric-mean florescence intensity (gMFI) in PbTII GFP^+^ cells and polyclonal naïve T cells. Bar graph shows the ratio of Tbet gMFI in PbTII cells relative to polyclonal naïve T cells. (A,B) Data pooled from 3 independent experiments for sgCD19, sgCXCR5, and sgCXCR3+sgCXCR5 cells and pooled from 2 independent experiments for sgCXCR3 cells (n= 5 mice per group, per independent experiment). (C) Representative FACS plots and bar graph of direct ex vivo staining of CCL5^+^ in sgCD19, sgCXCR5, or sgCXCR3+sgCXCR5 GFP^+^ PbTΙΙ cells at day 7 p.i. Data pooled from 2 independent experiments (n= 5 mice per group, per independent experiment). Bars indicate median. Statistical test performed using paired two-way ANOVA with Tukey’s multiple comparison test. p-values are indicated where *p< 0.05, **p< 0.01.

