## Supplementary Methods for "Spatial transcriptomics maps molecular and cellular requirements for CD4^+^ T cell-dependent immunity to malaria"

Supplementary Methods Figure 1

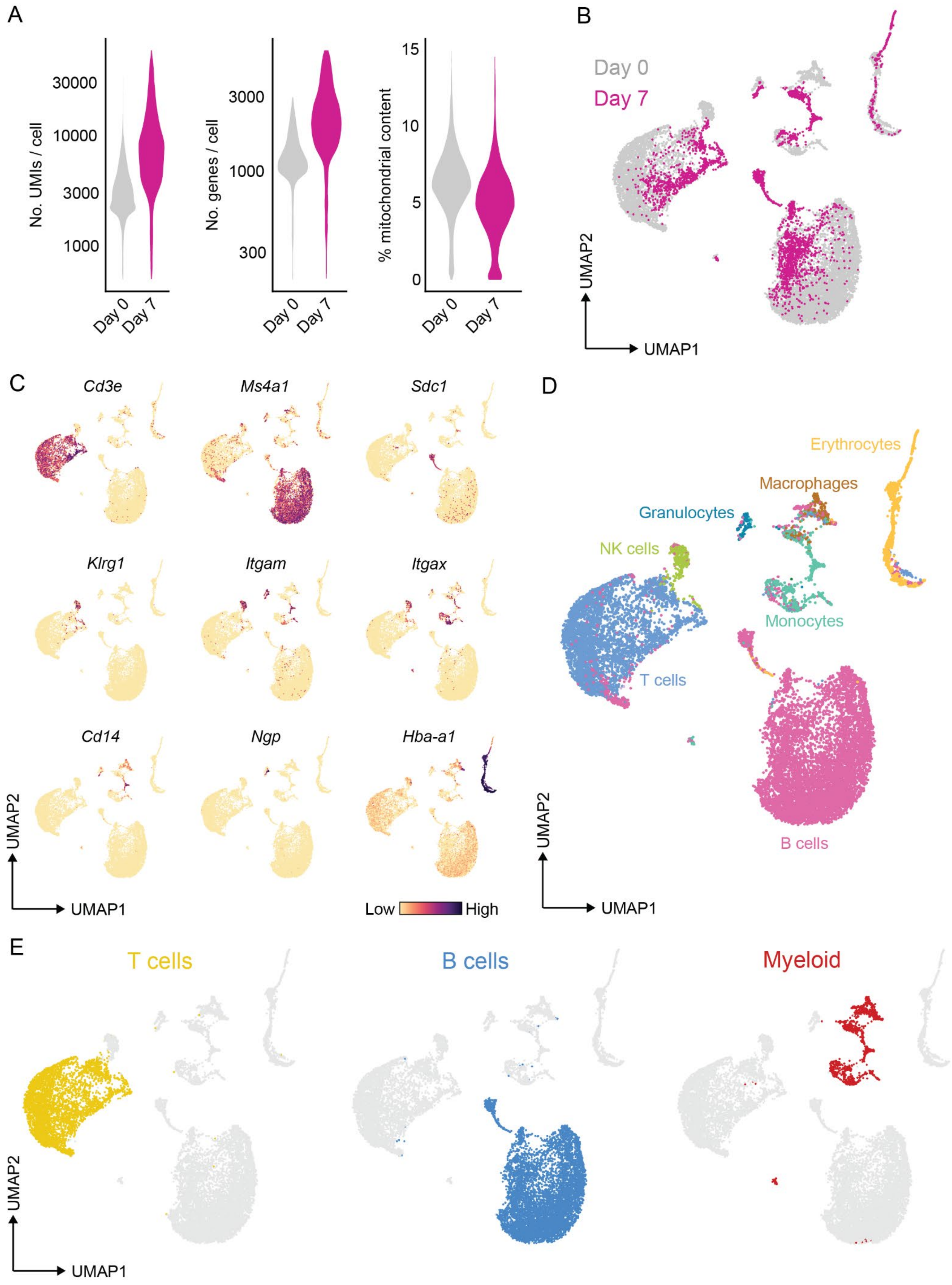

**Supplementary Methods Figure 1. scRNA-seq profiles unbiased splenic subsets before and during malaria infection.**

(A) QC metric assessment of single cell transcriptomes (no. UMIs/cell; no. Genes/cell and % mitochondrial content) post-filtering of low quality cells and doublets with DoubletDecon. (B) UMAP embedding of day 0 and day 7 single splenocyte transcriptomes. (C) Gene expression highlighting major splenocyte subsets: T cells (*Cd3e*), B cells (*Ms4a1*), plasmablasts (*Sdc1*), NK cells (*Klrg1*), monocytes and macrophages (*Itgam, Itgax, Cd14*), granulocytes (*Ngp*) and erythrocytes (*Hba-a1*). (D) Reference-based automatic cell type annotations generated with SingleR. (E) UMAP embeddings showing highlighted clusters identified as T cells (gold), B cells (blue), and myeloid cells (red) which were advanced for further analysis.

Supplementary Methods Figure 2

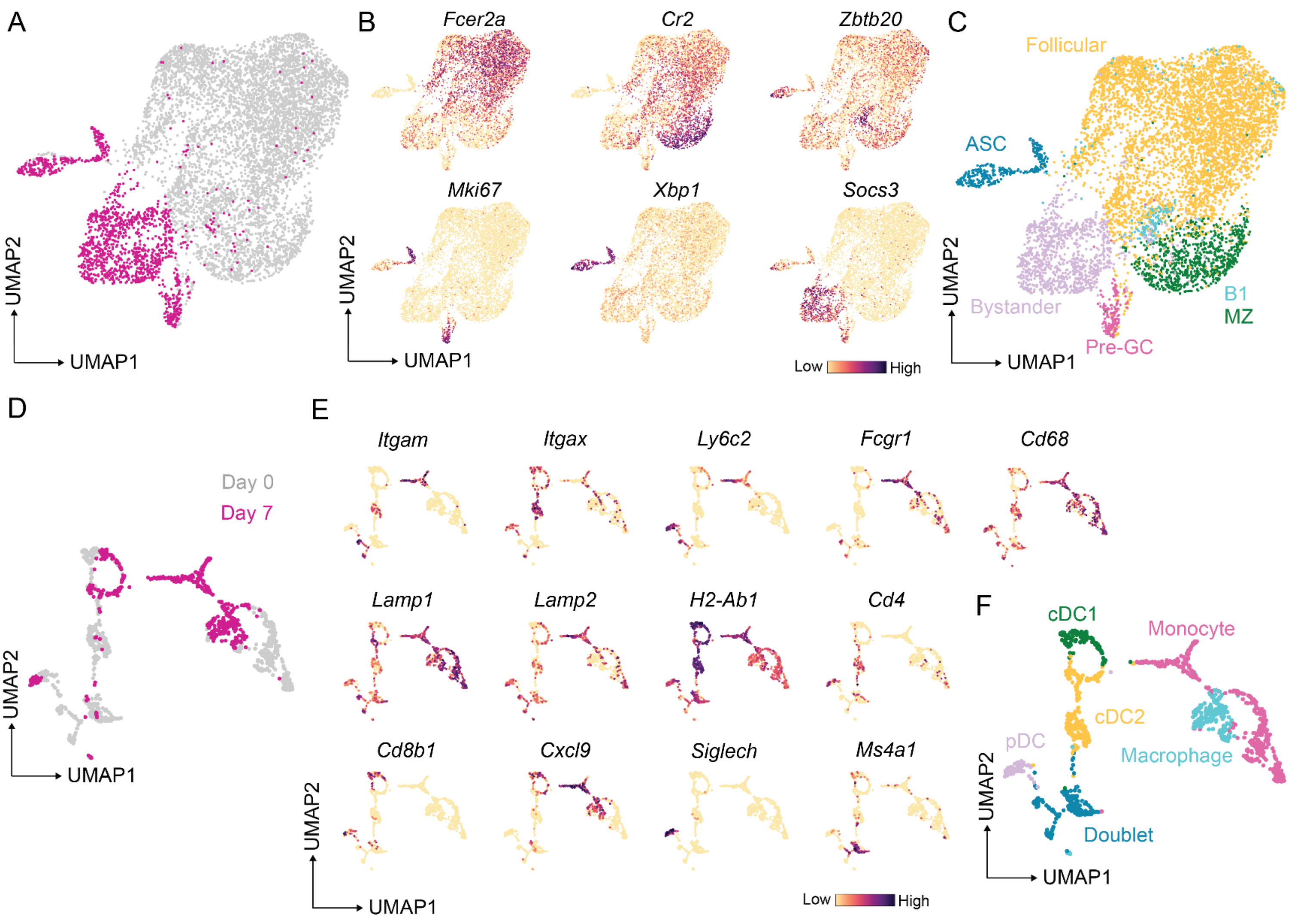


**Supplementary Methods Figure 2. Unbiased single cell transcriptomics of splenocytes reveals B cell and myeloid subsets.**

(A) UMAP embedding of B cells identified in Supp. Methods Fig. 1E, grouped according to collection before (day 0) or during (day 7) malaria infection.

(B) Gene expression revealing follicular (*Fcer2a*), bystander (*Socs3*), MZ (*Cr2*), B1 (*Zbtb20*), pre-GC (*Mki67*) and ASCs (*Xbp1*) B cell subsets.

(C) Inferred B cell subsets based on gene expression in (B).

(D) UMAP embedding of myeloid cells identified in Supp. Methods Fig. 1E, grouped according to collection before (day 0) or during (day 7) malarial infection.

(E) Gene expression revealing myeloid cell subsets.

(F) Inferred myeloid cell subsets based on gene expression in (E).

Supplementary Methods Figure 3

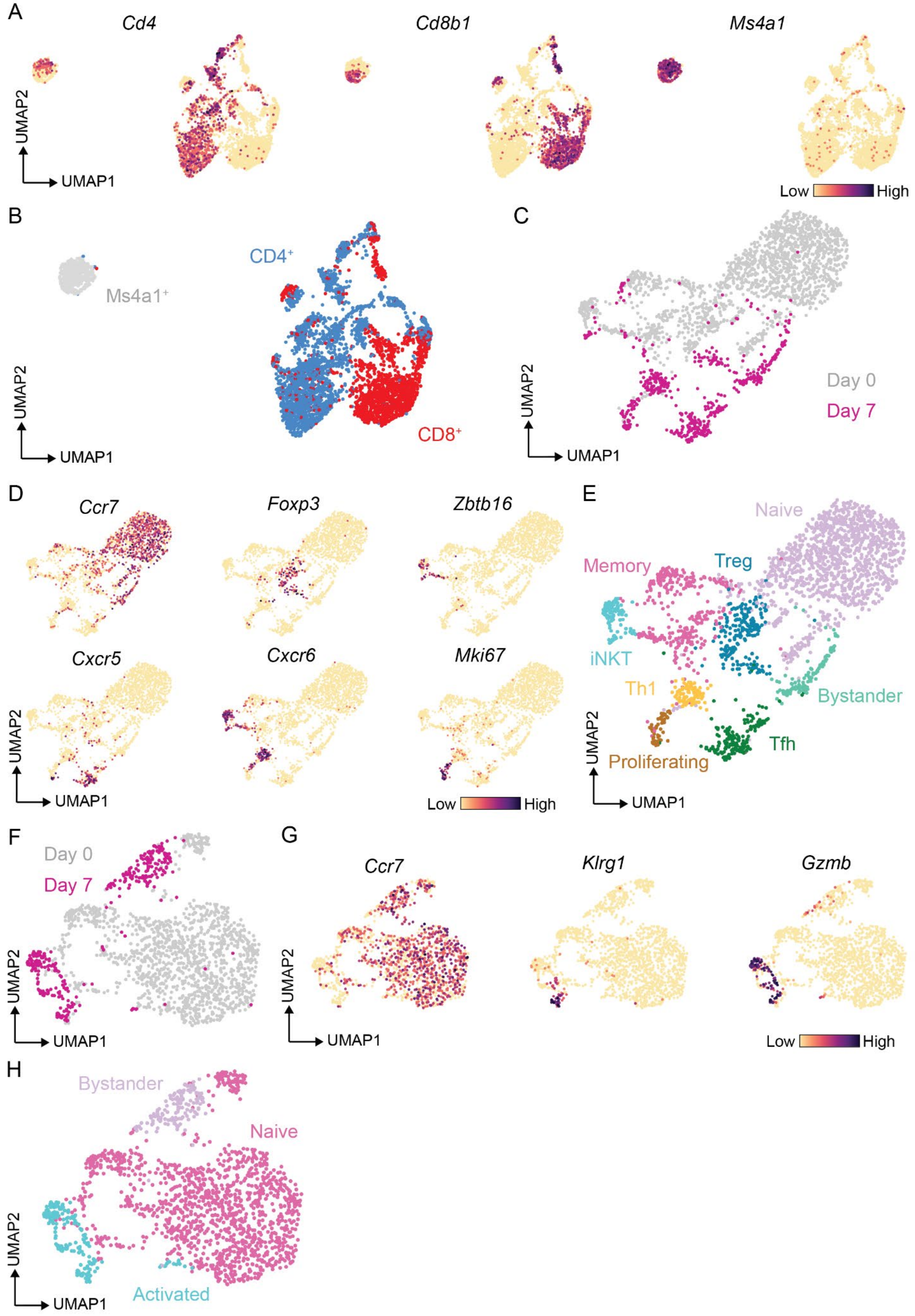

**Supplementary Methods Figure 3. Unbiased single cell transcriptomics of splenocytes reveals CD4$^{+}$ and CD8$^{+}$ T cell subsets.**

(A) Gene expression in UMAP embeddings of T cell transcriptomes showing *Cd4* and *Cd8b1,* identifying CD4$^{+}$ and CD8$^{+}$ T cells respectively. *Ms4a1* identifies doublets which were removed. (B) Classification of T cells as CD4$^{+}$, CD8$^{+}$, or as doublets based on *Ms4a1.* (C) UMAP embeddings of CD4$^{+}$ T cells grouped according to collection before (day 0) or during (day 7) malarial infection. (D) Gene expression revealing functional CD4$^{+}$ T cell subsets: naïve (*Ccr7*), Treg (*Foxp3*), iNKT (*Zbtb16, Cxcr6*), Tfh (*Cxcr5*), Th1 (*Cxcr6*) and proliferatinng cells (*Mki67*). (E) CD4$^{+}$ T cell subset annotations. (F) UMAP embeddings of CD8$^{+}$ T cells grouped according to collection before (day 0) or during (day 7) malarial infection. (G) Gene expression revealing functional CD8$^{+}$ T cell subsets: naïve (*Ccr7*) and activated T cells (*Klrg1, Gzmb*). (H) CD8$^{+}$ T cell subset annotations.
